# Analysis of HIV Reservoirs in Cellular Conjugates from Peripheral Blood

**DOI:** 10.1101/2021.01.19.427248

**Authors:** Liliana Pérez, Daniel Crespo-Vélez, Max Lee, Saami Zakaria, April Poole, Jennifer Bell, Stephen A. Migueles, Tae-Wook Chun, Susan Moir, Frank Maldarelli, Eli A. Boritz

**Affiliations:** Virus Persistence and Dynamics Section, Vaccine Research Center, National Institute of Allergy and Infectious Diseases, Bethesda, MD, USA; Laboratory of Immunoregulation, National Institute of Allergy and Infectious Diseases, Bethesda, MD, USA; Clinical Retrovirology Section, HIV Dynamics and Replication Program, National Cancer Institute, Frederick, MD, USA

**Keywords:** HIV Infections, CD4-Positive T-Lymphocytes, Humans, Flow Cytometry, Phylogeny, Polymerase Chain Reaction (PCR)

## Abstract

Defining distinctive attributes of HIV-infected cells will inform development of HIV cure-directed therapies. Prior *ex vivo* studies of blood and tissue have suggested that some HIV-infected CD4 T cells are found in conjugates with other cell types. Here, we analyzed levels and sequences of HIV nucleic acids in sorted cellular conjugates from PBMC. Compared to single CD4 T cells, conjugates containing CD4 T cells showed no enrichment for HIV DNA or RNA. However, in several HIV controllers, HIV DNA sequences from sorted conjugates were enriched for sequences closely related to plasma viruses. In ART-treated people, although subgenomic HIV DNA sequences in sorted conjugates and single cells were genetically intermingled, intact proviruses were more frequent in whole blood cells than in magnetically-purified CD4 T cells. We conclude that some HIV-infected cells have attributes that predict preferential loss during sample processing, and that may also reflect vulnerability to therapeutic targeting *in vivo*.

## Introduction

Defining attributes that may distinguish HIV-infected CD4 T cells from their uninfected counterparts *in vivo* is a major challenge in HIV cure-related research. Central to this challenge are the low frequency of HIV-infected cells during chronic infection and the lack of clear virus protein expression on many of these cells. Because HIV-infected cells account for a small minority of all CD4 T cells *ex vivo* (*Brenchley et al., 2004*), analysis of unfractionated CD4 T cells from these compartments largely reflects the attributes of uninfected cells. At the same time, although cell culture procedures that induce virus gene expression have enabled HIV-infected cell detection and sorting assays (*Baxter et al., 2016; Cohn et al., 2018*), HIV latency reversal *in vitro* can be inefficient (*Ho et al., 2013*) and can also change cellular attributes (*Pardons et al., 2019*). As a result, many research groups have used a hypothesis-driven “reverse” approach whereby cell-associated HIV is quantified and characterized within FACS-sorted CD4 T cell subsets. These studies have revealed higher levels of HIV DNA in cells sorted by maturation state (*Boritz et al., 2016; Brenchley et al., 2004; Chomont et al., 2009*), functional profile (*Gosselin et al., 2010*), and anatomic localization (*Yukl et al., 2010*). Qualitative differences between subsets have also been observed, including the presence and nature of intracellular HIV transcripts (*Telwatte et al., 2018*) and both culture- (*Banga et al., 2018; Fukazawa et al., 2015)* and sequence-based *(Kuo et al., 2020; Lee et al., 2017*) measures of replication competency. Nevertheless, HIV enrichment across CD4 T cell subsets in these studies has been modest, with a large majority of uninfected cells in all subsets sorted to date.

Complementing studies of physiologic CD4 T cell subsets as HIV hosts, several reports have examined relationships between “non-classical” CD4 T cell surface markers and intracellular HIV (*Abdel-Mohsen et al., 2018; Darcis et al., 2020; Descours et al., 2017; Hogan et al., 2018; Iglesias-Ussel et al., 2013; Raposo et al., 2017; Serra-Peinado et al., 2019; Vásquez et al., 2019*). The IgG2 receptor CD32A was identified as a specific marker of HIV-infected CD4 T cells in people receiving antiretroviral therapy (ART), with a proposed mechanism involving intracellular sensing of HIV and induction of a battery of non-CD4-T-cell genes (*Descours et al., 2017*). We and others were unable to confirm the marked HIV enrichment in CD32^+^ CD4 T cells, tempering hopes for emerging HIV cure strategies based on CD32 (*Badia et al., 2018; Bertagnolli et al., 2018; Osuna et al., 2018; Pérez et al., 2018*). Nevertheless, several studies did find higher levels of HIV RNA in CD32^+^ CD4 T cells (*Abdel-Mohsen et al., 2018; Vásquez et al., 2019*). As with CD32, the tumor marker CD30 and the B cell marker CD20 have been detected on a minority of HIV-infected CD4 T cells and associated with elevated levels of cell-associated HIV RNA (*Hogan et al., 2018; Serra-Peinado et al., 2019*). Although these findings do not immediately suggest means of completely eliminating HIV-infected cells, they could lead to novel approaches for suppressing an “active reservoir” as part of “functional cure” strategies.

In some cases, CD4 T cells bearing mixed lineage markers may arise through cellular conjugate formation. Although cell—cell conjugates can form routinely during PBMC processing even from uninfected people, the formation of conjugates containing CD4 T cells has been previously associated with HIV infection. Examples of this include the propensity of CD4 T cells from people with AIDS to bind to natural killer cell target cells and to monocytes (*Barnaba et al., 1988; Dudhane et al., 1996*). Conjugates between CD4 T cells and activated platelets are also more abundant in PBMC from people with HIV than in HIV-negative PBMC (*Green et al., 2015; Real et al., 2020; Simpson et al., 2020*). Moreover, we and others found that for the specific case of CD32, apparent surface expression by CD4 T cells in PBMC was often attributable to cellular conjugate formation (*Osuna et al., 2018; Pérez et al., 2018*). It is important to acknowledge that these studies did not rule out true CD32 expression on very rare CD4 T cells with some degree of enrichment for HIV DNA, as one more recent study has suggested (Darcis et al., 2020). Nevertheless, the unequivocal identification of rare cells showing atypical phenotypes in FACS data from heterogeneous blood and tissue samples is technically challenging. The relationship among CD32, intracellular HIV, and cell—cell conjugates also calls into question whether HIV-infected CD4 T cells, or a subpopulation of these cells, might preferentially form conjugates with other cells or cellular material, either *in vivo* or during sample processing.

Here we investigated enrichment for HIV in peripheral blood CD4 T cells showing mixed-lineage marker expression. We reasoned that some HIV-infected CD4 T cells might preferentially form conjugates with other cells or cellular material due expression of adhesion markers, virus proteins, or early cell death phenotypes that can be recognized by receptors on other cells. We examined this issue first in people with spontaneous control of HIV in the absence of ART (i.e., HIV controllers). We selected HIV controllers for the study because our previous work in this population revealed rare circulating CD4 T cells harboring HIV DNA with a recognizable genetic signature of recent infection (*Boritz et al., 2016*). Thus, we anticipated that enrichment for recently infected cells, potentially forming part of an “active reservoir,” might be most easily assessed by sequencing in controllers. Subsequently, we used a combination of technologies in an attempt to extend our findings in controllers to the setting of ART.

## Results

### FACS of cell—cell conjugates containing CD4 T cells in PBMC

We collected cellular conjugates from PBMC in people with HIV by using FACS to sort CD3^+^CD4^+^ material bearing mixed lineage markers. We used only freshly acquired samples with no previous cryopreservation for these experiments. Our upstream gating steps did not include area-by-height forward light scatter gating typically used for doublet exclusion, enabling collection of single CD4 T cells as well as three presumptive cell—cell conjugate populations based on expression of CD14 and CD32 (*Figure 1*). Cellular events with low forward light scatter and CD32 staining were considered as single CD4 T cells (*Figure 1*). Events with high forward light scatter that were CD3^+^CD4^+^CD32^−^ were considered to be presumptive conjugates, as were CD3^+^CD4^+^CD32^+^ events that were either negative or positive for CD14 (*Figure 1*). For simplicity these populations are identified in the figures as FSC^lo^, FSC^hi^, CD14^−^, and CD14^+^. Presumptive conjugate populations together accounted for ~10% of all cellular events from HIV+ PBMC, at levels that were similar between HIV controllers and ART-treated people (*Figure 1; Table I*).

**Figure 1.**
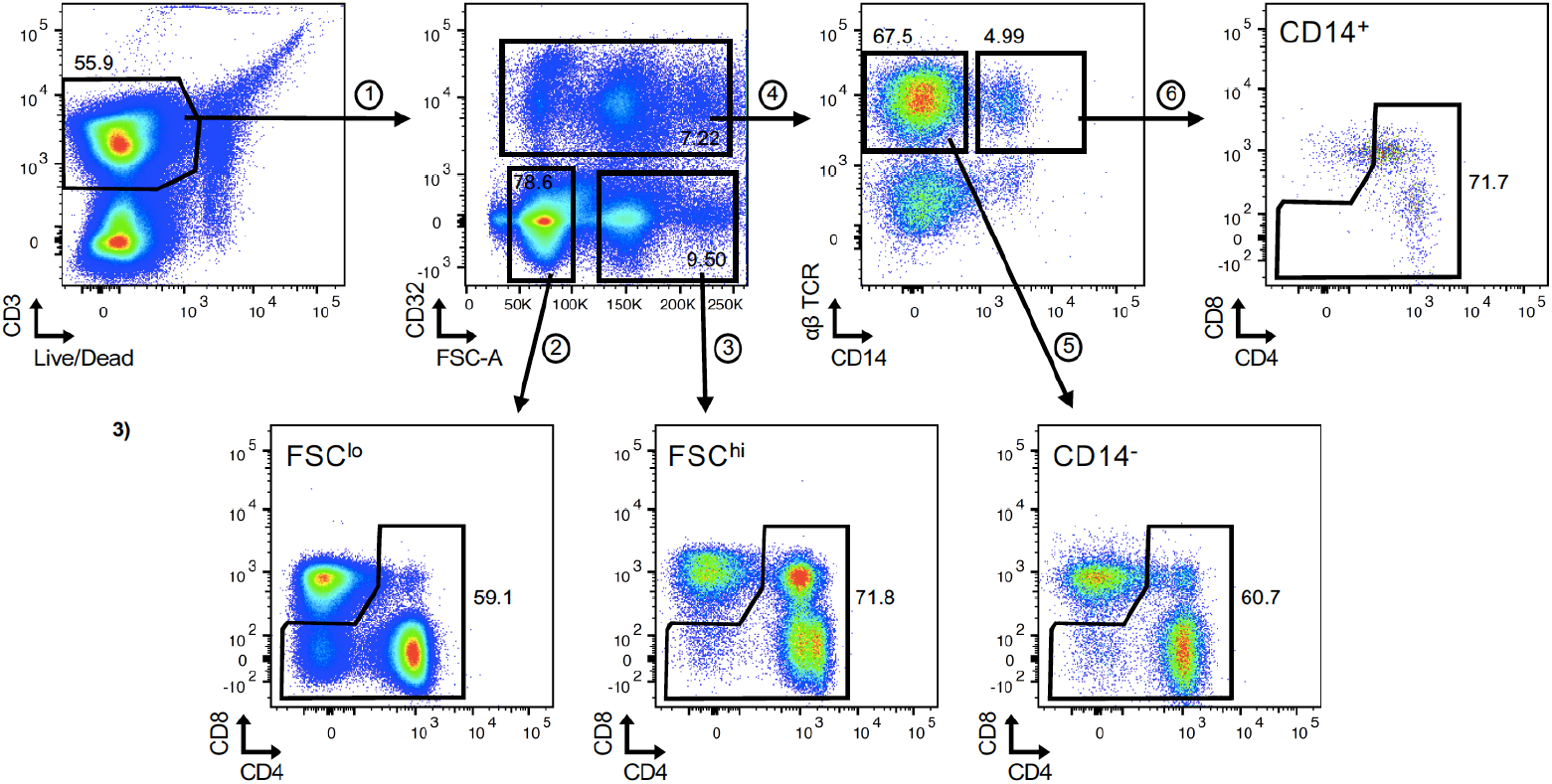
Flow cytometry of single CD4 T cells and CD4-T-cell–cell conjugates from PBMC. PBMC isolated from leukapheresis or whole blood that were CD3^+^ and viable were separated by (1) CD32 expression and forward light scatter. CD32^−^ cells were divided into two populations: (2) single CD4 T cells showing a low forward light scatter (FSC^lo^) and (3) high forward light scatter (FSC^hi^). CD32^+^ cells were gated based on their expression of (4) αβ TCR and CD14. Cells that were only positive for TCR αβ were further divided into (5) CD14^−^ and (6) CD14^+^. CD4^−^ and CD4^+^ T cells were sorted from these four cell populations: FSC^lo^, FSC^hi^, CD14^−^, and CD14^+^.

**Table I.**
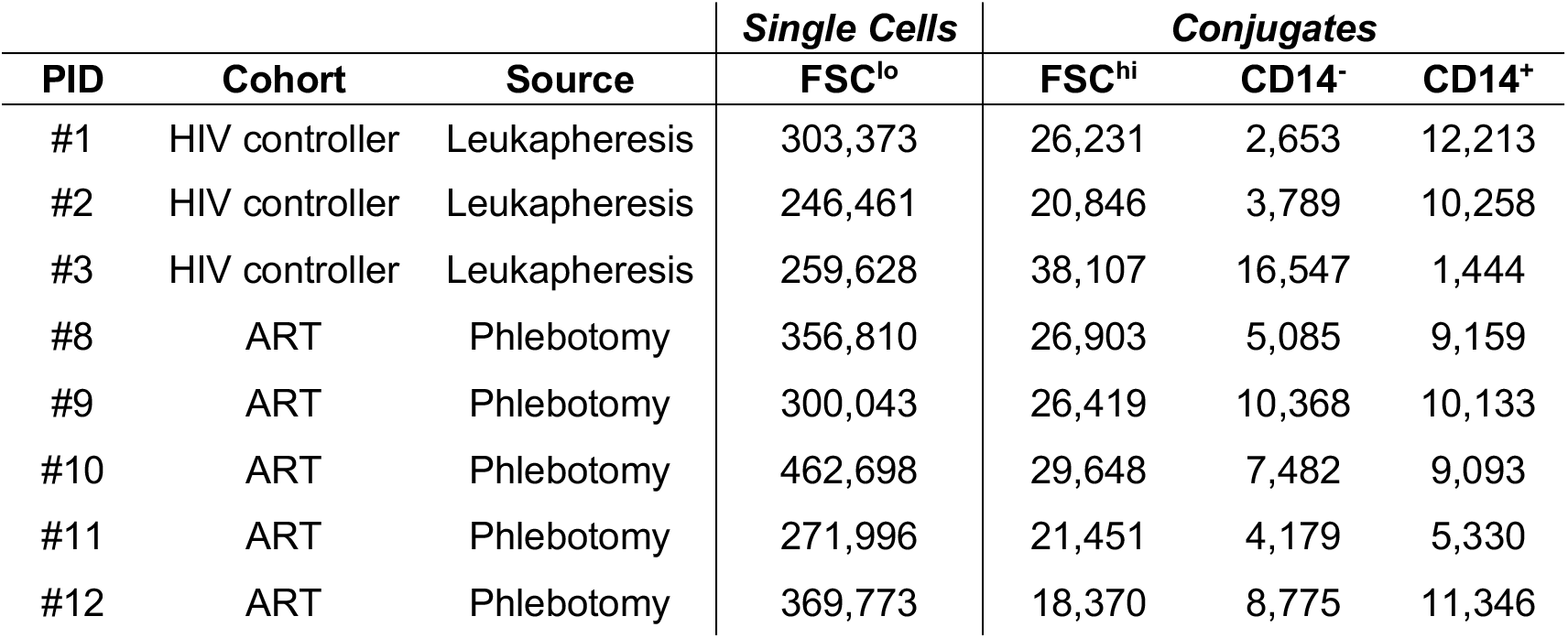
Number of FSC^lo^, FSC^hi^, CD14^−^ and CD14^+^ sorted events in HIV controllers and ART-treated participants.

To better characterize sorted cell populations as in our previous study of CD3^+^CD4^+^CD32^+^ material, we performed post-sort FACS analysis. As in our previous study, this showed heterogeneous patterns consistent with a predominance of true conjugates in FSC^hi^, CD14^−^, and CD14^+^ populations (*Pérez et al., 2018; Figure1—figure supplement 1*). In particular, the FSC^hi^ population showed a predominance of CD4 and CD8 T cells, suggesting T-cell—T-cell conjugates. The CD14^−^ population appeared on post-sort analysis as CD32^−^ T cells mixed predominantly with B cells. The CD14^+^ population appeared as CD32^−^ T cells mixed with monocytes. Small numbers of events retaining the mixed-lineage marker expression of the original sorted material were detected in CD14^−^ and CD14^+^ populations, consistent either with conjugates that failed to separate during sorting or with true single cells bearing mixed lineage markers. As expected, most sorted cells in the FSC^lo^ population were CD4 T cells. Overall, therefore, we were able to use FACS to enrich PBMC samples for conjugates between CD4 T cells and other cell types. These conjugates occurred at frequencies that were much higher than typical frequencies of HIV infection in PBMC *ex vivo.*

### Quantification of HIV nucleic acids in cell—cell conjugates

We quantified HIV DNA and RNA in these sorted populations both from HIV controllers and ART-treated participants. Using limiting dilution PCR for a fragment of the *env* gene, we detected no enrichment for HIV DNA in any of the three conjugate populations, as compared to single CD4 T cells. This was true in HIV controllers, where levels of HIV DNA were ~1/5,000 events or lower (*Figure 2A*), and in ART-treated people, where levels were approximately 10-fold higher (*Figure 2B*). We next used qRT-PCR to quantify unspliced and spliced transcripts in cell RNA from a subset of participants for whom extracted RNA was available. Levels of these transcripts were uniformly low in single CD4 T cells and all three conjugate populations both from HIV controllers and ART-treated people (*Figure 2C, D*). Therefore, HIV infection was detected both in single CD4 T cells and in cellular conjugates containing CD4 T cells from PBMC of HIV controllers and ART-treated people, with enrichment neither for HIV DNA nor for transcriptionally-active HIV in conjugate populations.

**Figure 2.**
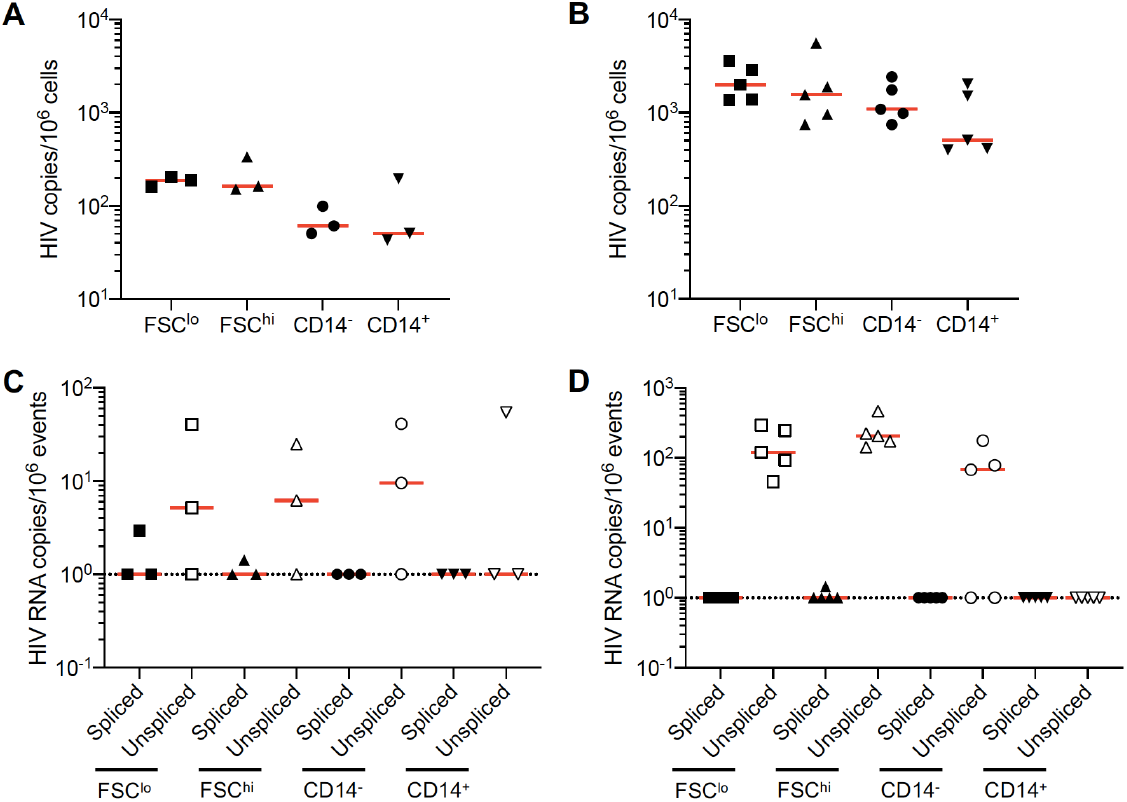
Levels of cell-associated HIV DNA and RNA in FSC^lo^, FSC^hi^, CD14^−^, and CD14^+^ cells, sorted from PBMC of HIV controllers and ART-treated participants. Copies of HIV DNA per million sorted FSC^lo^, FSC^hi^, CD14^−^, and CD14^+^ cells from PBMC isolated from (A) leukapheresis in HIV controllers or (B) phlebotomy in ART-treated participants. Copies of spliced (close) and unspliced (open) HIV RNAs in sorted events from PBMC of (C) HIV controllers and (D) ART-treated participants. To allow display of these wide-ranging values—including several values of zero—on a logarithmic scale, each plotted value represents the measured value + 1. In all figures, horizontal bars denote median values and dashed line indicates limit of detection.

### HIV DNA sequence analysis in cell—cell conjugates

In a previous study, we found that the HIV-infected CD4 T cell pool in blood from HIV controllers consisted predominantly of expanded cellular clones harboring archival proviruses, with a much smaller population of cells harboring viruses genetically similar to plasma viruses (*Boritz et al., 2016*). To determine in the present study whether such recent sequences might be preferentially found in conjugate populations, we performed Sanger sequencing and phylogenetic analysis on single-copy *env* PCR products from conjugates, single cells, and plasma virions in HIV controllers. In two of three participants, we found extensive HIV genetic intermingling both among sorted cell subsets and between each subset and plasma virions (*Figure 3A; participants #1 and #2*). In these two participants, there was no clear segregation between HIV DNA sequences in conjugates and those in single CD4 T cells. In the third participant, however, we found that while most HIV DNA sequences showed a clustered, archival pattern, rare sequences were genetically similar to plasma viruses (*Figure 3; participant #3*). These sequences were exclusively found in FSC^hi^ and CD14^−^ conjugate populations. In this individual, recent HIV DNA sequences in conjugate populations represented <3.5% of all HIV-infected cells in conjugates and <1% of all HIV DNA copies detected in blood cells (*Figure 3B*). Interestingly, analysis of a parallel sample from the same participant acquired by phlebotomy instead of leukapheresis did not uncover this rare population of HIV DNA sequences (*Figure 3—figure supplement 2; participant #3*). Overall, therefore, although conjugates were not enriched for HIV nucleic acids compared to single CD4 T cells, we found enrichment for very rare cells with markers of an actively replicating pool in conjugates from one HIV controller, as well as evidence that recovery of this cell population was influenced by upstream PBMC processing steps.

**Figure 3.**
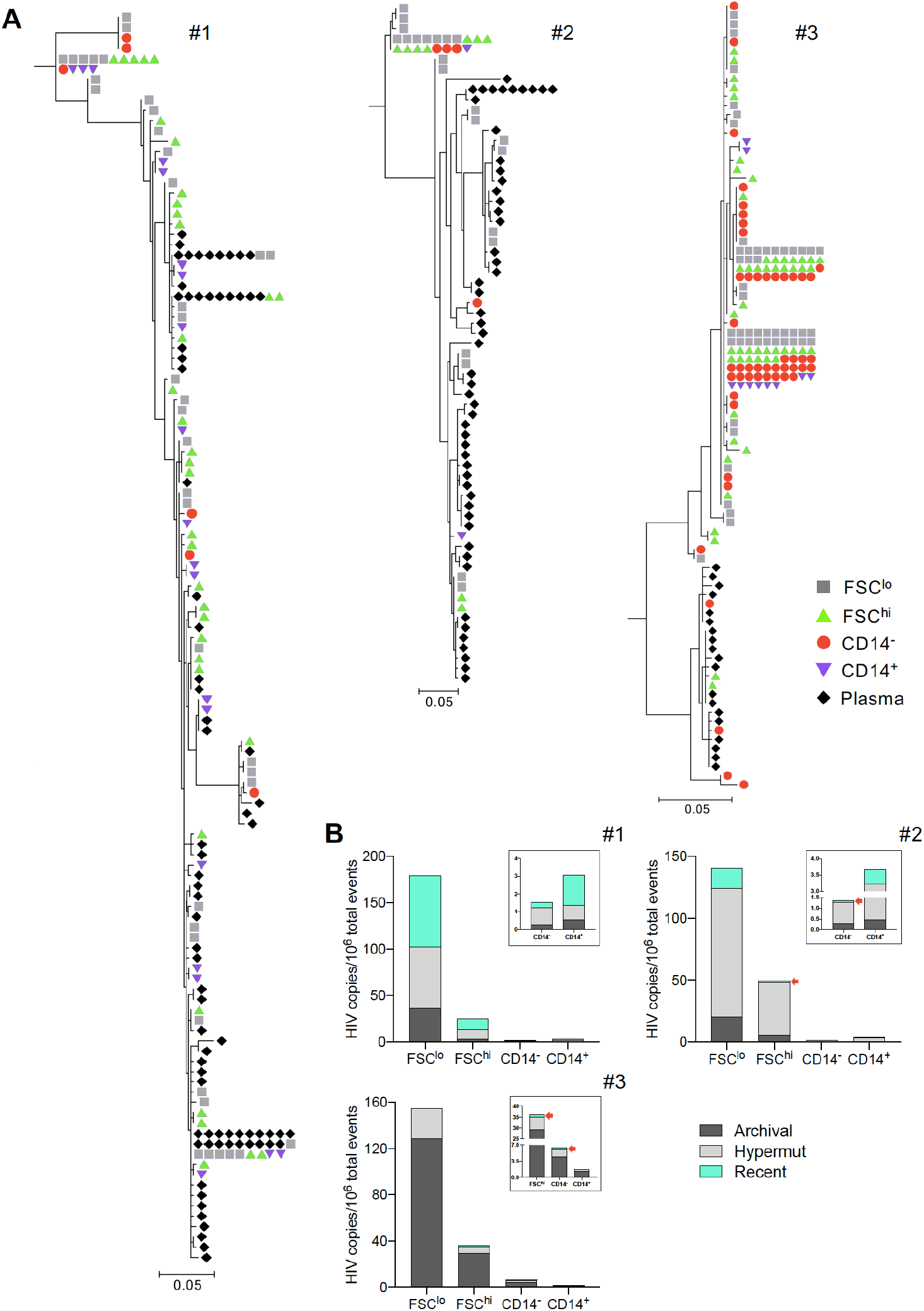
Subgenomic sequence analysis from FSC^lo^, FSC^hi^, CD14^−^, and CD14^+^ cells, sorted from leukapheresis PBMC of HIV controllers. (A) Sequences of individual HIV DNA copies were determined by Sanger sequencing of products obtained by fluorescence-assisted clonal amplification, which amplifies a region of the HIV *env* gene. Phylogenetic trees were constructed as described in the Methods. Gray squares represent sequences from FSC^lo^ cells, green triangles are sequences from FSC^hi^ cells, red circles are sequences from CD14^−^ cells, and purple downward facing triangles sequences from CD14^+^ cells. (B) HIV copies/10^6^ events displaying archival (dark gray), recent (light blue), and G-to-A hypermutated (Hypermut, light gray) HIV DNA detected in FSC^lo^, FSC^hi^, CD14^−^, and CD14^+^ cells. In some participants, red arrows also indicate the proportion of recent HIV DNA.

We also performed Sanger sequencing and phylogenetic analysis on single-copy *env* PCR products from conjugates and single cells in ART-treated participants. As in the three HIV controllers, we found that most HIV DNA sequences from conjugates in five ART-treated participants were genetically similar to HIV DNA sequences from single cells over the subgenomic region analyzed. Moreover, clusters of matching sequences representing possible expanded cellular clones invariably contained sequences both from single cells and from conjugates (*Figure 4A*). In all individuals, single CD4 T cells and conjugates contained similar levels of hypermutated HIV DNA (*<18% of all HIV DNA sequences; Figure 4B*). Thus, this approach did not suggest a distinct genetic subpopulation of HIV DNA sequences in cell—cell conjugates from ART-treated people.

**Figure 4.**
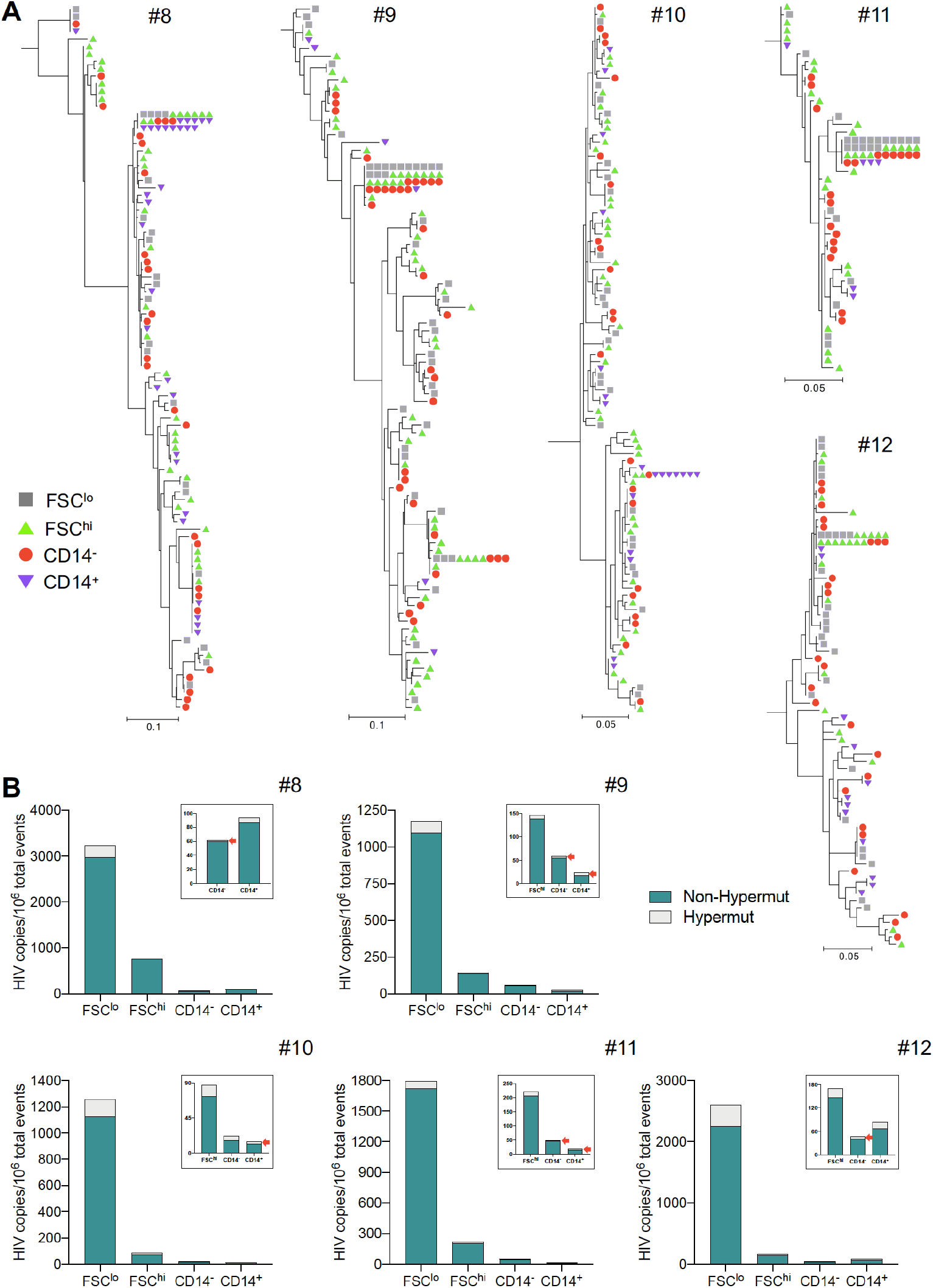
Subgenomic sequence analysis from FSC^lo^, FSC^hi^, CD14^−^, and CD14^+^ cells, sorted from whole blood PBMC of ART-treated participants. (A) Phylogenetic trees were constructed using sequences of individual HIV DNA copies determined by Sanger sequencing of products obtained by fluorescence-assisted clonal amplification. Gray squares represent sequences from FSC^lo^ cells, green triangles from FSC^hi^ cells, red circles from CD14^−^ cells, and purple downward facing triangles from CD14^+^ cells. (B) HIV copies/10^6^ events displaying non-hypermutated (Non-Hypermut, teal), and G-to-A hypermutated (Hypermut, light gray) HIV DNA detected in FSC^lo^, FSC^hi^, CD14^−^, and CD14^+^ cells. In some participants, red arrows also indicate the proportion of hypermutated HIV DNA.

### FACS of CD4-T-cell—platelet conjugates in PBMC

Based on these findings and on previous evidence of interaction between CD4 T cells and activated platelets in HIV infection (*Green et al., 2015; Real et al., 2020; Simpson et al., 2020*), we expanded the focus of these studies to include CD4-T-cell—platelet conjugates. We detected CD4 T cells bearing platelet markers in freshly-acquired PBMC by labeling for CD42b +/− CD62P, and employed a gating strategy in which CD4 T cell conjugates with other cells (Non-CD4^+^) were isolated first, followed by collection of CD4-T-cell—platelet conjugates (Plt^+^) and single CD4 T cells (*Figure 5*). Using this strategy, we found that CD4-T-cell—platelet conjugates accounted for 2.5-11.6% of all cellular events in PBMC from HIV+ participants (*Table II*). Post-sort analyses confirmed that while cell—cell conjugates contained mixtures of CD4 T cells and various cell types, CD4-T-cell—platelet conjugates contained predominantly CD4 T cells (*Figure 5—figure supplement 3*). Platelet markers were no longer detected on CD4 T cells after sorting in the platelet conjugate population, suggesting platelet dissociation during sorting.

**Figure 5.**
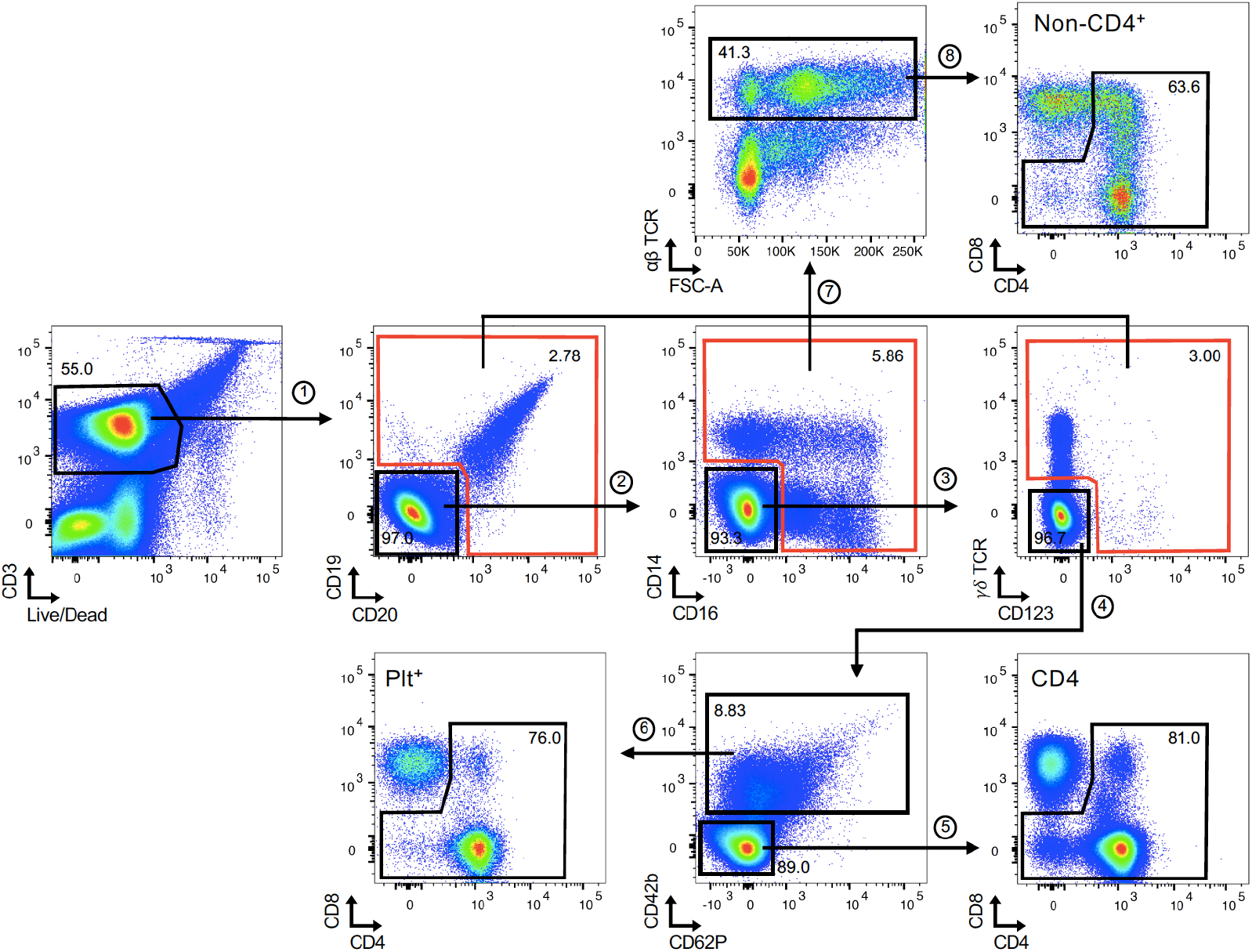
Flow cytometry of single CD4 T cells, CD4-T-cell–platelets, and CD4-T-cell–cell conjugates from PBMC. PBMC isolated from whole blood that were CD3^+^ and viable were gated by expression (1) CD19 and CD20, (2) CD14 and CD16, (3) γδ TCR and CD123. Cells that were CD19^−^, CD20^−^, CD14^−^, CD16^−^, γδ TCR^−^, and CD123^−^ were further separated by expression of platelet surface markers: (5) CD42b^−^ and CD62P^−^ (CD4) and (6) CD42b^+^ and CD62P^+/−^ (Plt^+^). Cells that were CD19^+^, CD20^+^, CD14^+^, CD16^+^, γδ TCR^+^, and CD123^+^ were (7) combined into one population and (8) αβ TCR^+^ cells were gated (Non-CD4^+^). CD4^−^ and CD4^+^ T cells were sorted from these three cell populations: CD4, Plt^+^, and Non-CD4^+^.

**Table II.** Number of CD4, Plt^+^, Non-CD4^+^ sorted events in HIV controllers and ART-treated participants.

| PID | Cohort | Source | Single Cells | Conjugates |  |
| --- | --- | --- | --- | --- | --- |
|  |  |  | CD4 | Plt<sup>+</sup> | Non-CD4<sup>+</sup> |
| #4 | HIV controller | Phlebotomy | 388,233 | 36,121 | 18,037 |
| #5 | HIV controller | Phlebotomy | 575,091 | 33,356 | 86,317 |
| #6 | HIV controller | Phlebotomy | 510,821 | 12,895 | 60,464 |
| #13 | ART | Phlebotomy | 369,815 | 28,341 | 26,413 |
| #14 | ART | Phlebotomy | 195,301 | 2,905 | 38,017 |
| #15 | ART | Phlebotomy | 540,498 | 62,640 | 58,306 |
| #16 | ART | Phlebotomy | 781,503 | 58,892 | 53,218 |

### Quantification of HIV DNA in CD4-T-cell—platelet conjugates

We next quantified HIV nucleic acids in CD4-T-cell—platelet conjugates in HIV controllers and ART-treated people. Again, using limiting-dilution PCR of a region of *env*, we found no enrichment for HIV DNA in CD4-T-cell—platelet conjugates or cell—cell conjugates, as compared to single CD4 T cells. HIV DNA levels were lower in HIV controllers (*~1 copy/10,000 cells; Figure 6A*) than in ART-treated people (*~1 copy/1,000 cells; Figure 6B*).

**Figure 6.**
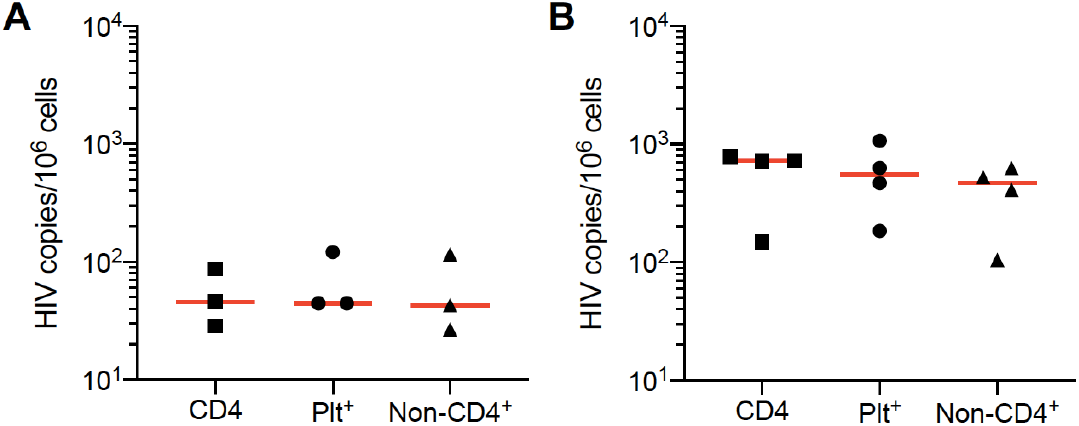
Levels of cell-associated HIV DNA in CD4, Plt^+^, and Non-CD4^+^ cells, sorted from PBMC of HIV controllers and ART-treated participants. Copies of HIV DNA per million sorted CD4, Plt^+^, and Non-CD4^+^ cells from PBMC isolated from whole blood in (A) HIV controllers and (B) ART-treated participants. In all figures, horizontal bars denote median values.

### HIV DNA sequence analysis in CD4-T-cell—platelet conjugates

We investigated genetic attributes of HIV sequences in CD4-T-cell—platelet conjugates by Sanger sequencing and phylogenetic analysis of single-copy *env* PCR products in these populations. In HIV controllers, we found an enrichment for HIV DNA sequences closely related to plasma virus sequences in CD4-T-cell—platelet conjugates and cell—cell conjugates in two of three participants (*Figure 7A; participants #4 and #5*). These HIV DNA sequences accounted for 36-60% of all HIV-infected cells in conjugates and 6.5-8% of all HIV DNA copies detected in blood cells (*Figure 7B*). The third HIV controller studied showed extensive HIV genetic intermingling between cell DNA and plasma virion populations, and also showed no clear segregation between HIV DNA sequences in conjugates and those in single CD4 T cells (*Figure 7A; participant #6*). In ART-treated participants, we found that most HIV DNA sequences from CD4-T-cell—platelet conjugates were genetically similar to HIV DNA sequences from single cells over the subgenomic region analyzed (*Figure 8A*). Clusters of matching sequences representing possible expanded cellular clones again included sequences both from single cells and from conjugates (*Figure 8A; participants #13 and #14*). The proportion of hypermutated HIV DNA sequences in all infected cells ranged from 10-68% in CD4 T cells and CD4-T-cell—cell conjugates (*Figure 8B*). Thus, as with cell—cell conjugates containing CD4 T cells, CD4-T-cell—platelet conjugates showed no overall enrichment for HIV DNA in HIV+ participants but did show enrichment for rare HIV DNA sequences with a genetic signature of recent infection in some HIV controllers studied.

**Figure 7.**
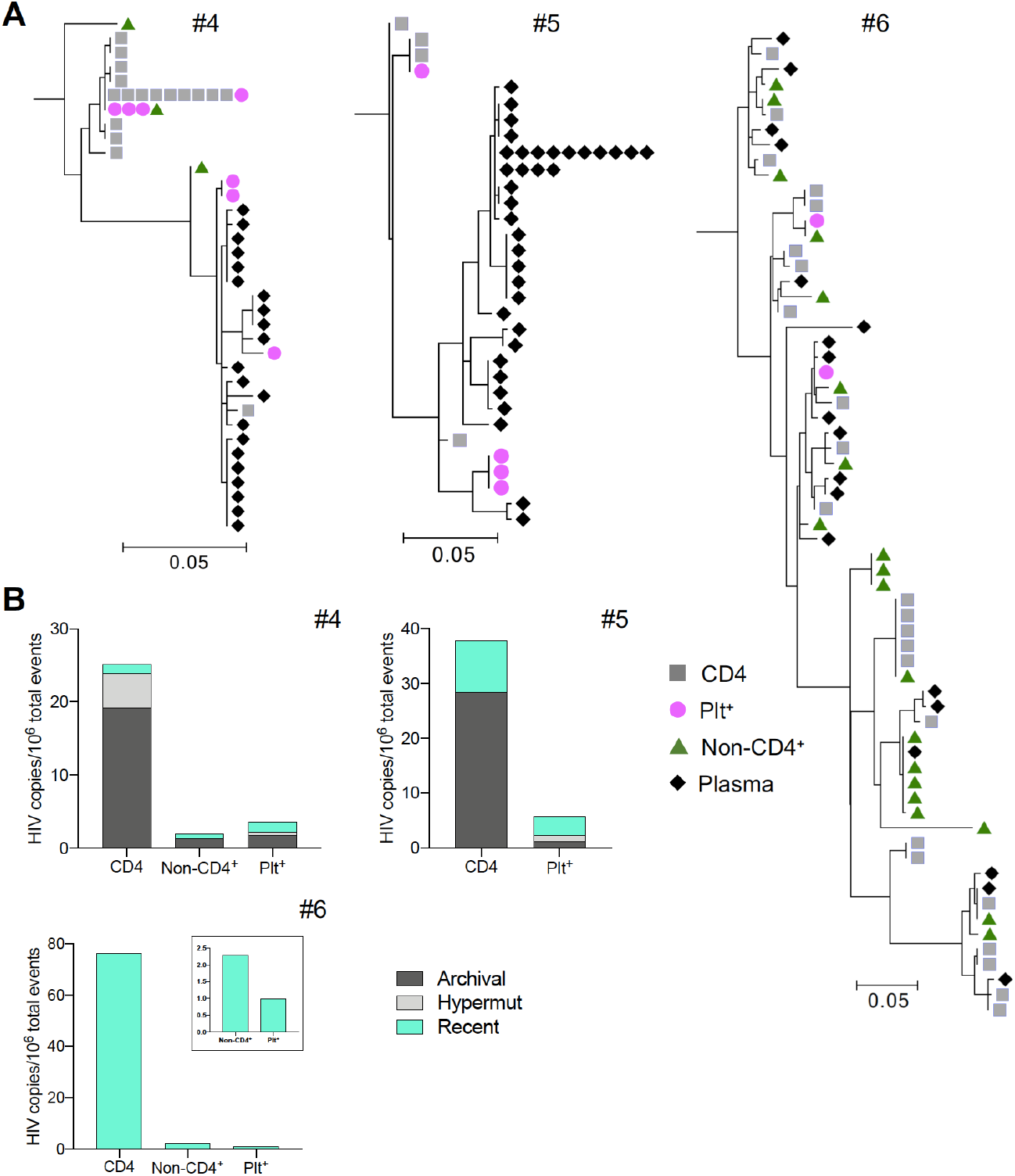
Subgenomic sequence analysis from CD4, Plt^+^, and Non-CD4^+^ cells, sorted from whole blood PBMC of HIV controllers. (A) Phylogenetic trees were constructed using sequences of individual HIV DNA copies determined by Sanger sequencing of products obtained by fluorescence-assisted clonal amplification. Gray squares represent sequences from CD4 T cells, pink circles from Plt^+^ cells, and dark green triangles from Non-CD4^+^ cells. (B) HIV copies/10^6^ events displaying archival (dark gray), recent (light blue), and G-to-A hypermutated (Hypermut, light gray) HIV DNA detected in CD4, Plt^+^ and Non-CD4^+^ cells.

**Figure 8.**
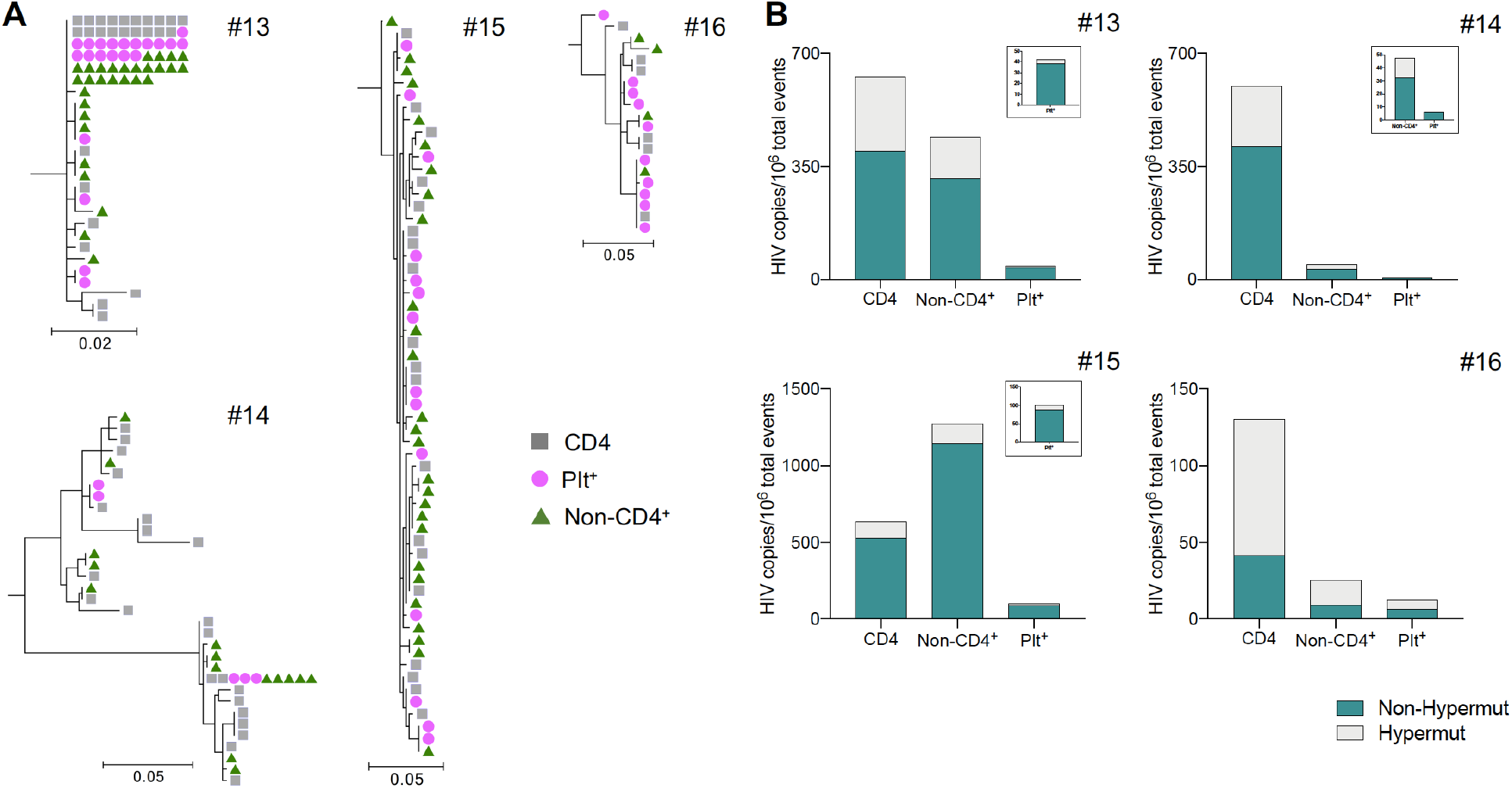
Subgenomic sequence analysis from CD4, Plt^+^, and Non-CD4^+^ cells, sorted from whole blood PBMC of ART-treated participants. (A) Phylogenetic trees were constructed using sequences of individual HIV DNA copies determined by Sanger sequencing of products obtained by fluorescence-assisted clonal amplification. Gray squares represent sequences from CD4 T cells, pink circles from Plt^+^ cells, and dark green triangles from Non-CD4^+^ cells. (B) HIV copies/10^6^ events displaying non-hypermutated (Non-Hypermut, teal), and G-to-A hypermutated (Hypermut, light gray) HIV DNA detected in CD4, Plt^+^ and Non-CD4^+^ cells.

### Genetically intact HIV DNA in whole blood vs. magnetically-purified CD4 T cells under ART

We next sought to extend positive findings in HIV controllers to ART-treated people. We noted important limitations in the above experiments, including the potential for preferential loss of labile cells during sample handling, and uncertainty about whether subgenomic virus sequencing can identify important subpopulations of HIV-infected cells during ART. Therefore, we sought to compare samples processed using standard procedures with samples subjected to minimal processing. We prepared peripheral blood from 10 ART-treated people by conventional density gradient centrifugation and magnetic CD4 T cell purification, and processed parallel aliquots of these samples by immediate whole blood cell DNA extraction. We then characterized HIV DNA in these samples using both single-copy *env* fragment sequencing and the intact proviral DNA assay (*IPDA; Figure 9A*), which simultaneously interrogates two regions of each virus DNA genome using multiplexed digital-droplet PCR (*Bruner et al., 2019*). As shown in Figure 9B, HIV *env* was detected by limiting dilution PCR in both whole blood and magnetically-purified CD4 T cell DNA. As expected, given the lack of enrichment for CD4 T cells in whole blood DNA, levels of HIV were lower in whole blood DNA samples. Single-copy sequence analysis showed evidence neither for sample contamination nor for distinct HIV genetic subpopulations in whole blood compared to CD4 T cells (*Figure 9—figure supplement 4*). Nonetheless, percentages of intact proviruses were significantly higher in whole blood cell DNA than in purified CD4 T cell DNA. This was true for all 8/10 participants in whom IPDA detected intact proviruses (*Figure 9C*). Overall, therefore, using a distinct experimental approach in samples from ART-treated individuals, we found evidence that a subset of cells containing intact HIV proviruses may be lost in standard CD4 T cell isolation procedures.

**Figure 9.**
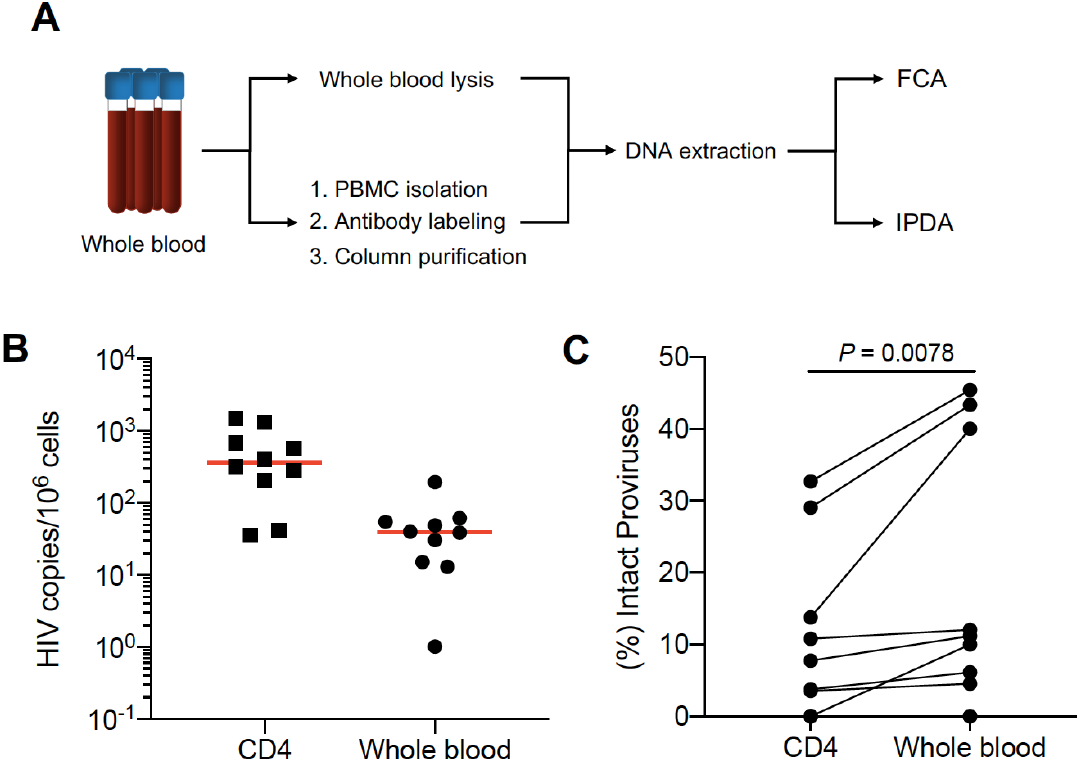
Levels of cell-associated HIV DNA and percentage of intact proviruses from whole blood and CD4 T cells of ART-treated participants. DNA was extracted from whole blood samples and negatively selected CD4 T cells of 10 ART-treated participants. DNA extracted from whole blood samples or magnetically-purified CD4 T cells from PBMC was assayed to measure (A) copies of cell-associated HIV DNA per million cells and (B) levels of intact, 3’ defective, and 5’ defective proviral HIV DNA by IPDA. The percent intact provirus was calculated by dividing the number of intact provirus by the total number of provirus HIV DNA detected in the sample. Horizontal bars denote median values. In (B) Wilcoxon, paired *t*-test p value is shown.

## Discussion

In this study, we characterized blood cell-associated HIV DNA genomes not recovered by conventional labeling and sorting procedures for single CD4 T cells. We began by quantifying and sequencing HIV DNA in FACS-sorted cell conjugates. This approach was informative in HIV controllers, with an enrichment in cell conjugates for a subset of HIV genomes closely resembling plasma viruses. These findings indicated that CD4 T cells harboring HIV sequences closely related to the actively replicating virus pool in HIV controllers may tend to adhere to other cell material and platelets either *in vivo* or during sample processing. In ART-treated people, by contrast, our initial FACS and subgenomic sequencing results did not reveal HIV genetic differences between conjugates and single CD4 T cells. Therefore, we extended our findings to the ART-treated setting by using IPDA to compare frequencies of intact HIV genomes between whole blood cells and magnetically-purified CD4 T cells. Our findings with this approach indicated that blood cells harboring genetically intact HIV may be preferentially lost during PBMC isolation, cell labeling, and negative selection of CD4 T cells using magnetic beads. Taken together, our results in this study suggest that subsets of HIV-infected blood cells harboring distinctive virus sequences also possess distinctive cellular attributes that can influence cell recovery during standard processing procedures.

We propose that our findings in HIV controllers and ART-treated people could have arisen from similar biological processes in these two clinical subgroups of PLWH. In HIV controllers, we have previously suggested genetic proximity between cell-associated HIV sequences and plasma viruses as a marker of recently infected cells (*Boritz et al., 2016*). Recently infected CD4 T cells might adhere to other cell material or platelets by expressing adhesion markers that relate to cellular activation, functional specialization, or early cell death, and that also relate to the tendency of certain CD4 T cell subtypes to become infected *in vivo*. Alternatively, our findings in HIV controllers may reflect direct effects of early, low-level virus gene expression on cellular adhesion, with recent infection serving as a genetic marker of enrichment for an “active reservoir”. In ART-treated people, by contrast, the persistence of ongoing HIV replication and thus the existence of recently infected cells are controversial (*Günthard et al., 1999; Josefsson et al., 2013; Kearney et al., 2014; Kieffer et al., 2004; Lorenzo-Redondo et al., 2016; Mens et al., 2007; Persaud et al., 2004; Ruff et al., 2002*). Nevertheless, loss of cells from the “intact reservoir” during CD4 T cell negative selection from PBMC would be expected if these cells expressed adhesion markers or virus proteins mediating conjugate formation. The prospect that active and intact reservoirs might overlap in ART-treated people – and that the cells of these reservoirs might be recognizable to other cells through interactions at the cell surface – warrants further study.

Although we applied rigorous FACS and molecular techniques to the samples in this study, our findings have important limitations. In FACS experiments, it was not possible to distinguish “physiologic” conjugates from those that may have formed during sample processing. We consider it likely that many conjugates in PBMC formed *in vitro*, as discussed previously for gastrointestinal-associated lymphoid tissue cells (*Thornhill et al., 2019*). Conjugates forming *in vitro* might depend on weak, non-specific interactions that permit dissociation during sorting and that are also not specific for cells of the active or intact HIV reservoirs. These considerations formed part of the rationale for our use of the alternative, whole blood cell lysis approach in samples from ART-treated people. A major challenge in this latter approach was the need to lyse cells and extract DNA in the absence of sorting or cell phenotyping information. Thus, it was not possible to confirm the nature of the cells harboring intact HIV genomes in this part of the study. It is even formally possible that some intact HIV DNA sequences in these experiments came from non-CD4 T cells, though previous studies do not suggest large intact HIV reservoirs in other circulating lymphoid or myeloid cell subsets (*Cattin et al., 2019; Durand et al., 2012; Josefsson et al., 2012; Massanella et al., 2019*). New technologies for single-cell imaging combined with *in situ* detection of HIV nucleic acids (*Baxter et al., 2016; Deleage et al., 2016; Grau-Expósito et al., 2017*), perhaps using early cell fixation or preservation to ensure recovery of labile cells, may be able to address these important issues.

In conclusion, our results raise key questions about the attributes of HIV-infected CD4 T cells that make up important subsets of HIV-infected reservoirs *in vivo*. It is important to emphasize that our results do not exclude the existence of rare HIV-infected cells expressing “non-classical” surface markers without conjugate formation, as has been demonstrated for CD20. Nonetheless, future studies should consider the potential for uneven cell losses during processing that could preferentially affect recently-infected cells or those containing genetically-intact HIV. Elucidating the mechanisms by which some of these cells may be recognized at the cell surface may help inform strategies for targeting HIV-infected cellular reservoirs or suppressing virus rebound upon ART interruption.

## Materials and Methods

### Fluorescence-Activated Cell Sorting (FACS)

Participant recruitment and informed consent were performed under IRB-approved protocols at NIH. For experiments evaluating CD4 T cell-conjugate populations (*Figures 1–4, Figure 1—figure supplement 1, and Figure 3—figure supplement 2*), freshly isolated PBMC obtained by leukapheresis or phlebotomy were stained with viability dye and the following fluorescently-labeled mAbs: CD3-APC-H7, CD4-BV785, CD8-PacBlue, CD14-BV650, CD16-PerCP/Cy5.5, CD19-BV605, CD20-BV570, CD27-Alx700, CD32-PE, CD123-PE/Cy5, CD45RO-ECD, TRC αβ-FITC, and TCR γδ-APC. PBMC were then sorted without doublet exclusion on a BD FACSAria into four populations based on the expression of CD32 and CD14: 1) CD32^−^and low forward light scatter cells (FSC^lo^) (CD3^+^CD32^−^FSC^lo^CD4^+/−^), 2) CD32^−^ and high forward light scatter cells (FSC^hi^) (CD3^+^CD32^−^FSC^hi^CD4^+/−^), 3) CD32^+^CD14^−^ cells (CD14^−^) (CD3^+^CD32^+^TCRαβ^+^CD14^−^CD4^+/−^), and 4) CD32^+^CD14^+^ cells (CD14^+^) (CD3^+^CD32^+^TCRαβ^+^CD14^+^CD4^+/−^) (*Figure 1*). To characterize the composition of these cells, a portion of each population was re-analyzed on the flow cytometer after sorting (*Figure 1—figure supplement 1*).

In additional experiments evaluating CD4-T-cell–platelet conjugates (*Figures 5–8 and Figure 5—figure supplement 3*), whole blood obtained by phlebotomy was collected in tubes containing heparin as the anticoagulant. Freshly isolated PBMC were stained with viability dye and the following fluorescently-labeled mAbs: CD3-APC-H7, CD4-BV785, CD8-PacBlue, CD14-BV650, CD16-PerCP/Cy5.5, CD19-BV605, CD20-BV570, CD27-Alx700, CD32-PE, CD42b-BV711, CD62P- PE/Cy7, CD123-PE/Cy5, CD45RO-ECD, TRC αβ-FITC, and TCR γδ-APC. PBMC were then sorted without doublet exclusion into three populations: 1) single CD4 T cells (CD3^+^CD19^−^CD20^−^CD14^−^CD16^−^TCRγδ^−^CD123^−^CD42b^−^CD62P^−^CD4^+/−^), 2) CD4-T-cell–platelet conjugates (Plt^+^) (CD3^+^CD19^−^CD20^−^CD14^−^CD16^−^TCRγδ^−^CD123^−^CD42b^+^CD62P^+/−^CD4^+/−^), and 3) CD4-T-cell–cell conjugates (Non-CD4^+^) (CD3^+^CD19^+^CD20^+^CD14^+^CD16^+^TCRγδ^+^CD123^+^TCRαβ^+^CD4^+/−^) (*Figure 5*). To evaluate the composition of these cells, a portion of each population was re-analyzed on the flow cytometer after sorting (*Figure 5—figure supplement 3*).

### Extraction of cell-associated RNA and DNA

Cell conjugates were sorted by FACS into heat-inactivated fetal calf serum and kept on ice until further processing. Cells were then sedimented by centrifugation at 400 × *g* for 7 minutes at 4°C, lysed in RNAzol RT at <5 × 10^6^ cells/mL, homogenized by pipetting, and stored at −80°C until extraction. For RNA and DNA extraction, 0.4 volume of sterile H_2_O was added to each lysate to allow aqueous and organic phase separation. Total RNA was extracted from the aqueous phase according to the manufacturer’s instructions. The organic phase of each lysate was solubilized in DNAzol (Molecular Research Centers) to allow DNA extraction according to the manufacturer’s instructions.

### HIV DNA Quantification

Fluorescence-assisted clonal amplification (FCA) for single-copy quantification and sequencing of HIV DNA was performed as described (*Boritz et al., 2016*). Cell-associated HIV DNA levels were calculated as follows: (HIV DNA copies/10^6^ cells) = 10^6^ × (number of HIV DNA copies detected in FCA) ÷ (number of cell genome equivalents analyzed by FCA, as measured by quantitative albumin gene PCR). False-positive wells were defined as those that showed positive fluorescence traces in FCA but in which the presence of HIV *env* DNA could not be confirmed by nested PCR amplification and Sanger sequencing.

Archival, recent, non-hypermutated or hypermutated HIV DNA in blood belonging to each sorted population detected in HIV controllers or ART-treated participants were calculated as follows: (HIV copies/10^6^ events = No. of archival, recent, non-hypermutated or hypermutated sequences in population ÷ sum of all sequences in population) × 10^6^ × (sorted events in population ÷ sum of all sorted events in all populations) × (HIV DNA copies in population).

### HIV sequence analysis

A maximum of forty positive FCA products from each sample were further amplified by nested PCR and subjected to Sanger sequencing. Base calls for Sanger sequence reads were edited as previously described (*Boritz et al., 2016*). Edited sequences from all study participants were aligned using GeneCutter at www.hiv.lanl.gov. The alignment was improved manually and used for maximum-likelihood phylogenetic tree construction in MEGA, allowing identification and removal of rare contaminants. Sequences were then realigned by participant and used to create maximum-likelihood trees in DIVEIN (https://indra.mullins.microbiol.washington.edu/DIVEIN/) (*Deng et al., 2010*). Possible G-to-A hypermutated sequences were determined using analysis previously described (*Boritz et al., 2016*) and were excluded from the phylogenetic trees of each participant.

### Quantification of HIV RNA

Total RNA samples extracted from cell conjugates were treated with DNAse I for 20 minutes at 37°C and then 15 minutes at 70°C to inactivate the DNAse. Cellular total RNA samples were tested for HIV RNA by qRT-PCR for unspliced (*gag*) RNA or transcripts spliced between the SD1 and SA4 sites using previously described protocols (*Boritz et al., 2016*). Copies of RNA were enumerated using standard curves generated from dilutions of synthetic RNAs.

### Reverse transcription of HIV virion RNA

Virion RNA was extracted from pelleted virions of whole plasma samples using the QiaAmp vRNA mini kit (QIAGEN), according to the manufacturer’s instructions. Virion RNA samples for FCA and sequencing were denatured at 65°C for 10 minutes, annealed to *env* reverse transcription primer (envB3out) at a primer concentration of 100 nM, and reverse transcribed using SuperScript III (Life Technologies) at 50°C for 50 minutes followed by incubation at 85°C for 10 minutes to inactivate the reverse transcriptase.

### Extraction of DNA from Whole Blood and Magnetically-Purified CD4 Cells

In order to compare differences during sample preparation, whole blood samples obtained by phlebotomy were processed using two different methodologies. Whole blood was lysed using the Gentra Puregene Blood Kit (Qiagen). Also, PBMC were separated on a Ficoll gradient to isolate CD4 T cells by negative selection using a CD4 T cell isolation kit (Miltenyi Biotech). CD4 T cells were lysed using the cell lysis solution from the Gentra Puregene Blood Kit. DNA from whole blood samples and isolated CD4 T cells was extracted according to the manufacturer’s instructions.

### Intact Proviral DNA Assay (IPDA)

To quantify the intact, 3’ defective, and 5’ defective proviral HIV DNA, IPDA was perfomed on DNA extracted from whole blood samples and CD4 T cells isolated from PBMC by Accelevir Diagnostics. Description of this procedure has been previously published (*Bruner et al., 2019*). Results are shown as percent intact proviruses.

## Author Contributions

Conceptualization: L.P. and E.A.B. Data generation and analysis: L.P., M.L., D.C.V., and S.J.Z. Participant cohort and sample management: A.P., J.B., S.Mi., T-W.C., S. Mo., and F.M. Manuscript preparation: L.P. and E.A.B.

## Competing interests

The authors declare no competing interests.

## Accession Numbers

All DNA sequences in this manuscript have been deposited in GenBank under accession numbers XX-XXXX-XX-XXXX.

## Supplemental Figures

**Figure supplement 1.**
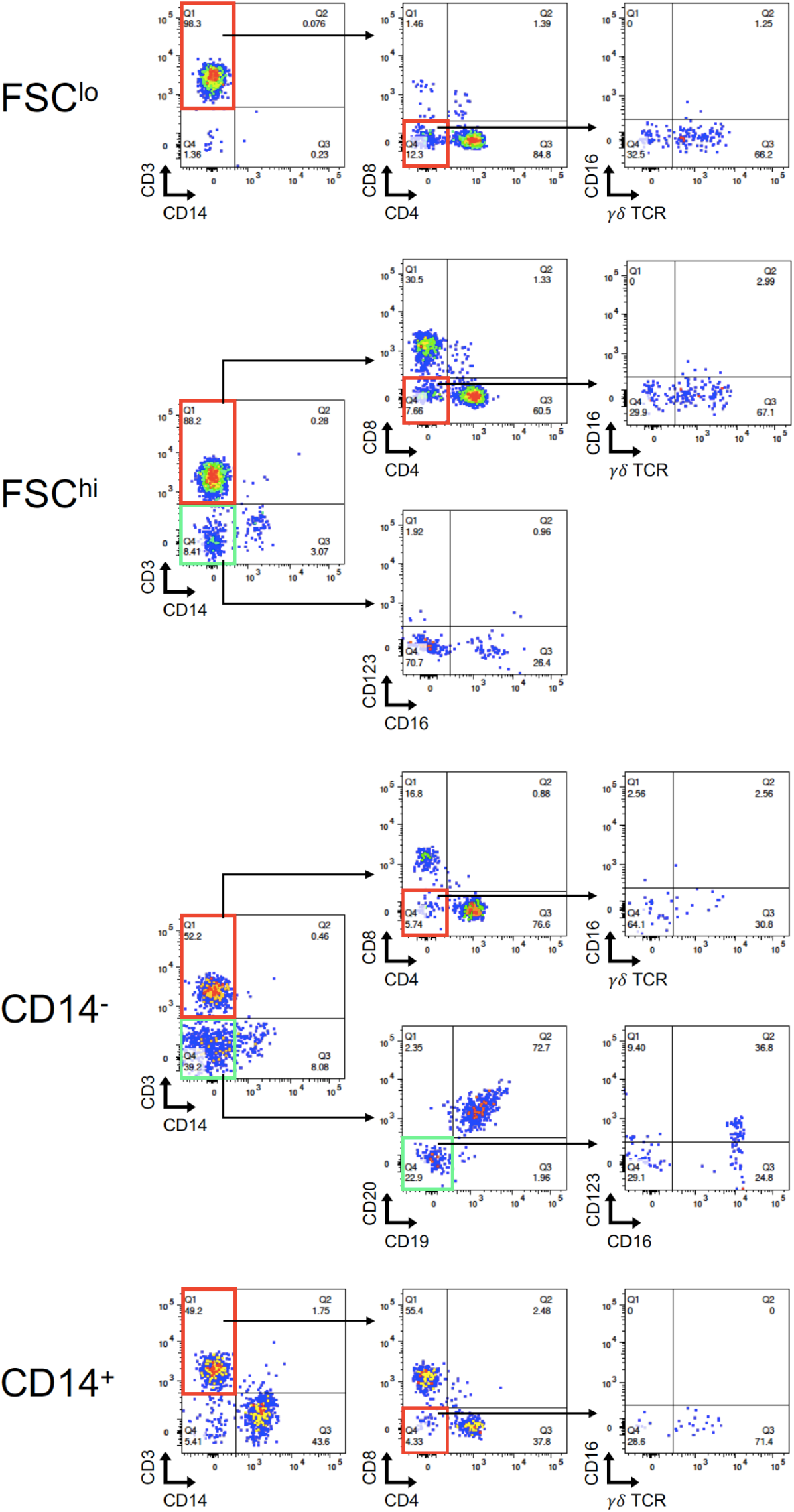
Post-sort flow cytometry of FSC^lo^, FSC^hi^, CD14^−^, and CD14^+^ cell populations. Cells were sorted as in Figure 1. FSC^hi^, CD14^−^, and CD14^+^ post-sorts revealed a heterogeneous pattern in conjugate populations. FSC^hi^ cells consists mainly of CD4-T-cell–CD8-T-cell conjugates, CD14^−^ cells of CD4 T cells mixed with B cells, and CD14^+^ cells of a mixture of CD4 T cells with monocytes.

**Figure supplement 2.**
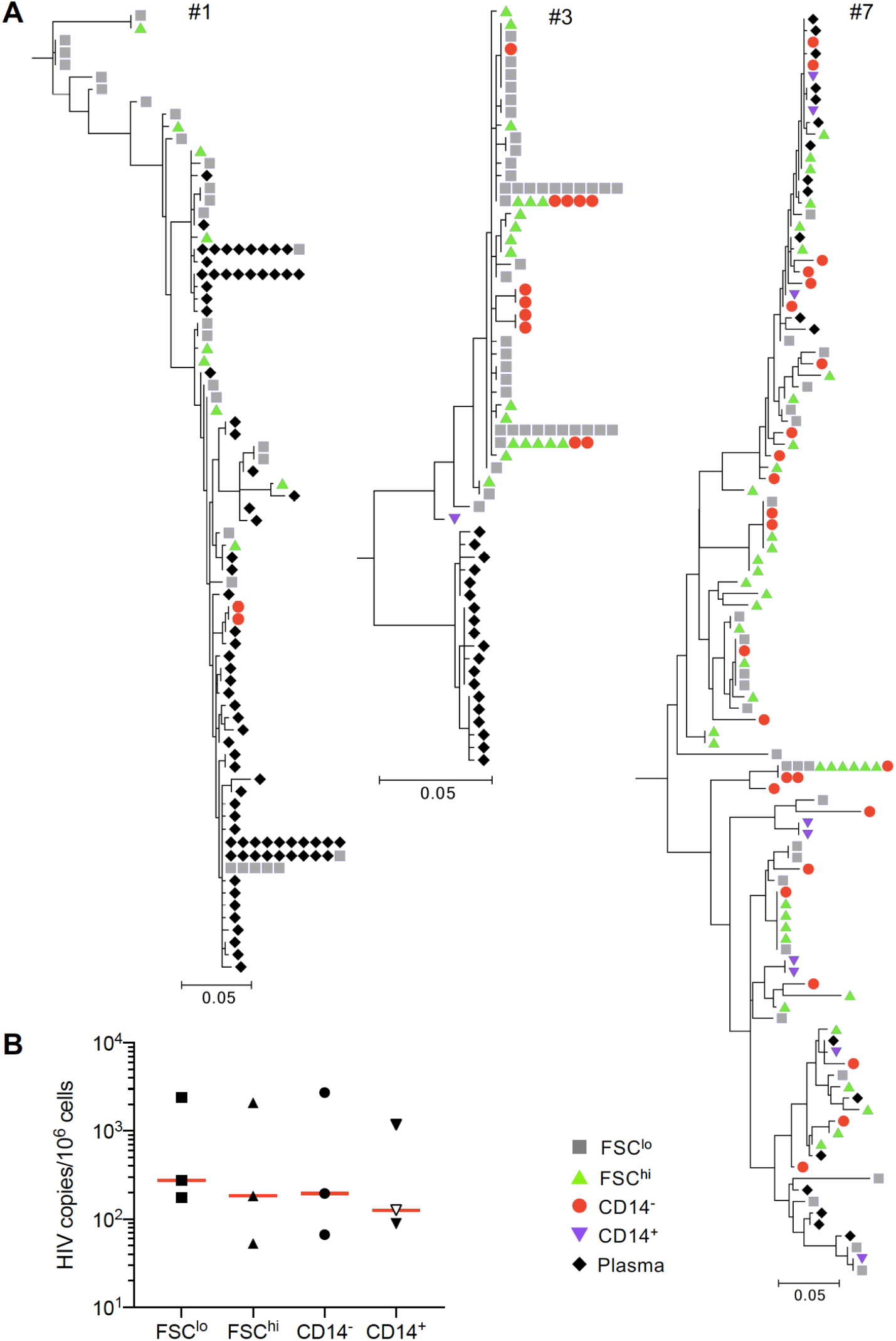
Subgenomic sequence analysis and levels of cell-associated HIV DNA from FSC^lo^, FSC^hi^, CD14^−^, and CD14^+^ cell populations sorted from whole blood PBMC of HIV controllers. (A) Sequences of individual HIV DNA copies were determined by Sanger sequencing of products obtained by fluorescence-assisted clonal amplification, which amplifies a region of the HIV *env* gene. Phylogenetic trees were constructed as described in the Methods. Gray squares represent sequences from FSC^lo^ cells, green triangles from FSC^hi^ cells, red circles from CD14^−^ cells, and purple downward facing triangles from CD14^+^ cells. (B) Copies of HIV DNA per million sorted FSC^lo^, FSC^hi^, CD14^−^, and CD14^+^ cells from PBMC isolated from whole blood in HIV controllers. Horizontal bars denote median values.

**Figure supplement 3.**
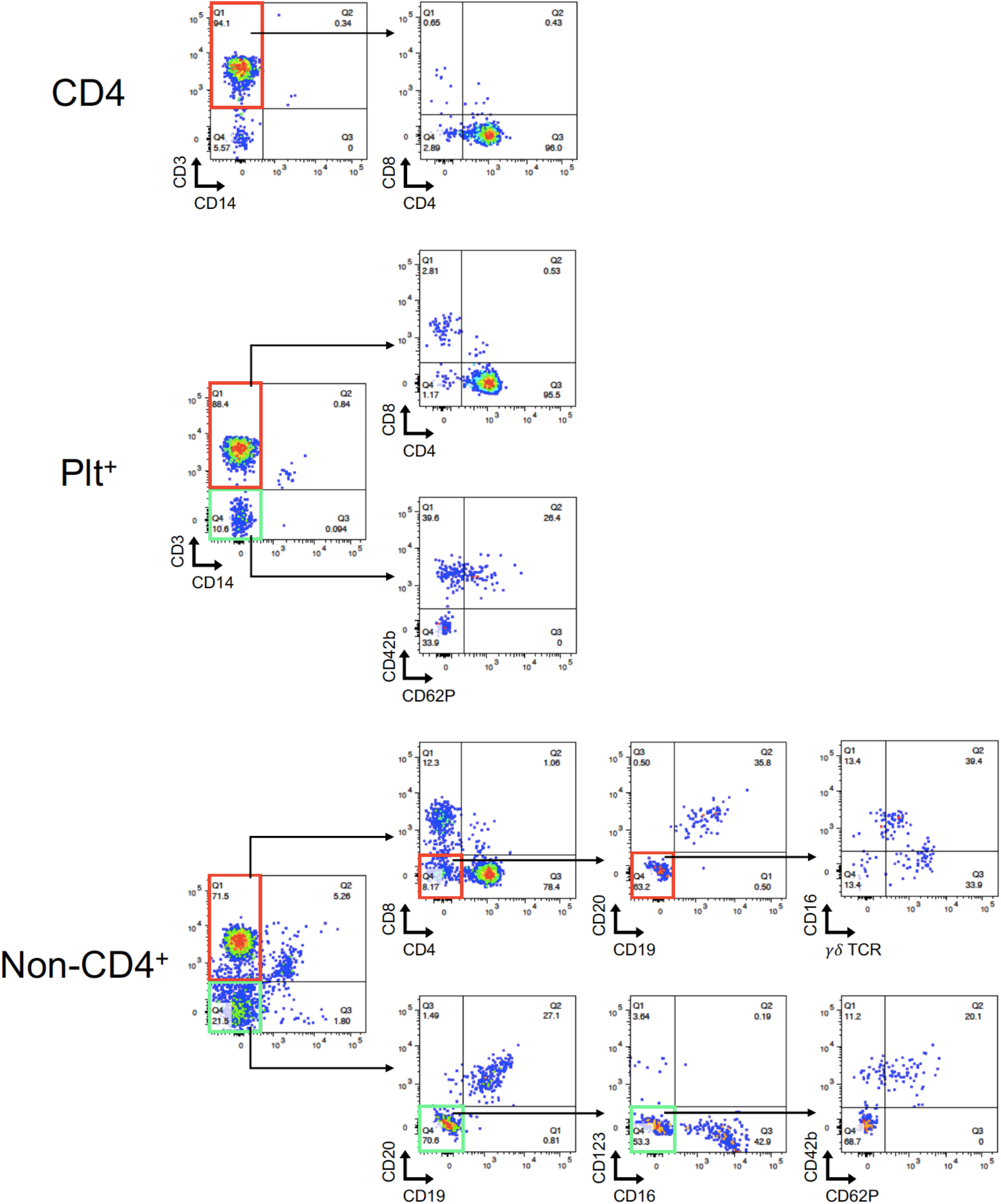
Post-sort flow cytometry of CD4, Plt^+^, Non-CD4^+^ cell populations. Cells were sorted as in Figure 5. Post-sorts analysis revealed a heterogeneous pattern in Non-CD4^+^ subset that consists of a mixture of CD4 T cells with various lineage markers. In contrast, the cells sorted based on platelet surface markers loses CD42b and CD62P expression on CD4 T cells.

**Figure supplement 4.**
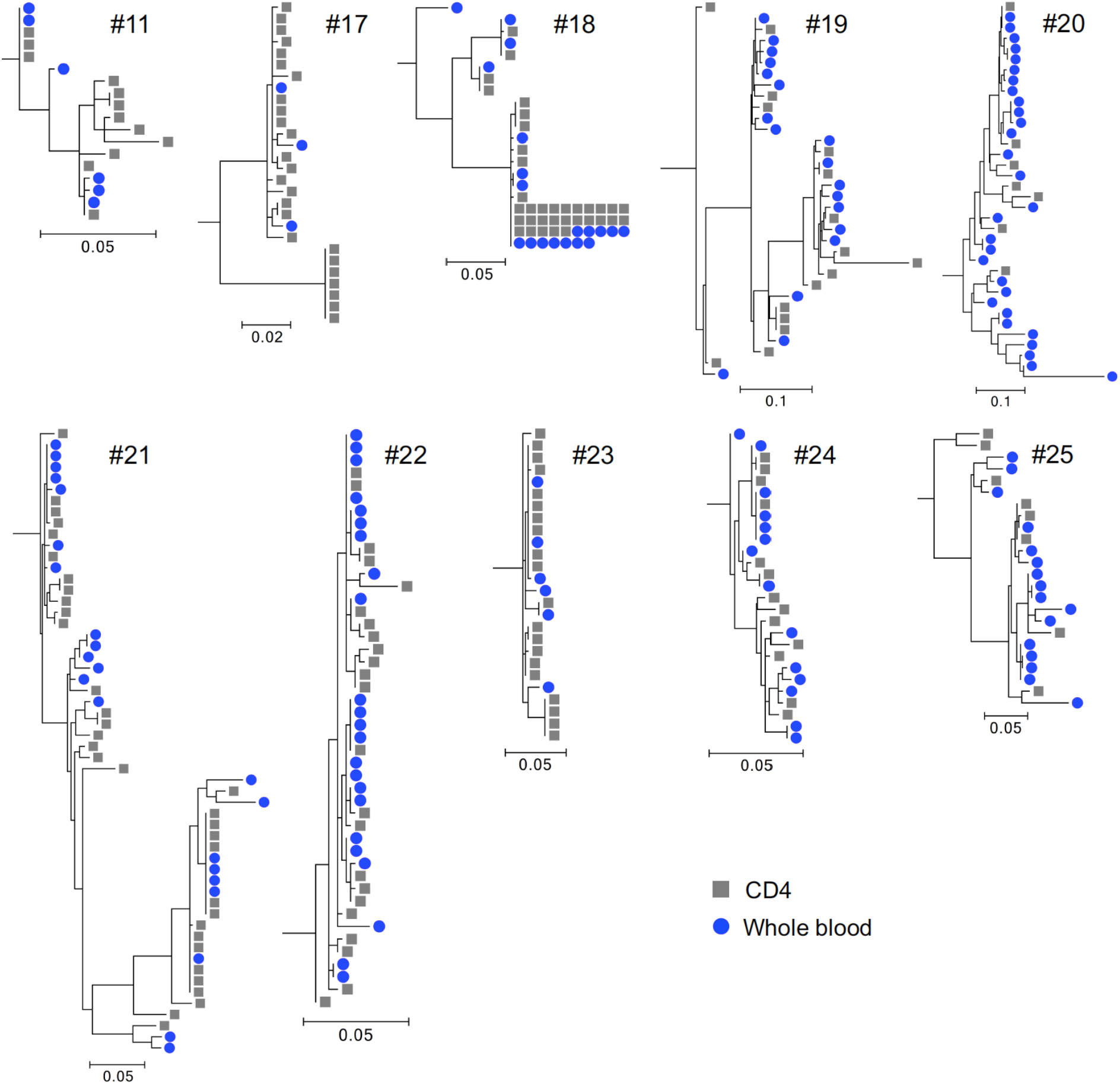
Subgenomic sequence analysis from whole blood and magnetically-purified CD4 T cells. Phylogenetic trees were constructed using sequences of individual HIV DNA copies determined by Sanger sequencing of products obtained by fluorescence-assisted clonal amplification. Gray squares represent sequences obtained from CD4 T cells and blue circles from whole blood.

## References

Abdel-Mohsen M, L Kuri-Cervantes, J Grau-Exposito, AM Spivak, RA Nell, C Tomescu, SK Vadrevu, LB Giron, C Serra-Peinado, M Genescà, J Castellví, G Wu, PM Del Rio Estrada, M González-Navarro, K Lynn, CT King, S Vemula, K Cox, Y Wan, Q Li, K Mounzer, J Kostman, I Frank, M Paiardini, D Hazuda, G Reyes-Terán, D Richman, B Howell, P Tebas, J Martinez-Picado, V Planelles, MJ Buzon, MR Betts, and LJ Montaner. 2018. CD32 is expressed on cells with transcriptionally active HIV but does not enrich for HIV DNA in resting T cells. Sci Transl Med 10. https://doi.org/10.1126/scitranslmed.aar6759, PMID: 29669853

Badia R, E Ballana, M Castellví, E García-Vidal, M Pujantell, B Clotet, JG Prado, J Puig, MA Martínez, E Riveira-Muñoz, and JA Esté. 2018. CD32 expression is associated to T-cell activation and is not a marker of the HIV-1 reservoir. Nat Commun 9: 2739. https://doi.org/10.1038/s41467-018-05157-w, PMID: 30013105

Banga R, FA Procopio, A Ruggiero, A Noto, K Ohmiti, M Cavassini, JM Corpataux, WA Paxton, G Pollakis, and M Perreau. 2018. Blood CXCR3(+) CD4 T Cells Are Enriched in Inducible Replication Competent HIV in Aviremic Antiretroviral Therapy-Treated Individuals. Front Immunol 9: 144. https://doi.org/10.3389/fimmu.2018.00144, PMID: 29459864

Barnaba V, G Valesini, G Gattamelata, R Benvenuto, A Velardi, and F Balsano. 1988. Increased number of CD4 cells able to bind to natural killer cell targets in the peripheral blood of AIDS related complex patients. Eur J Cancer Clin Oncol 24: 369–76. https://doi.org/10.1016/s0277-5379(98)90005-0, PMID: 2968260

Baxter AE, J Niessl, R Fromentin, J Richard, F Porichis, R Charlebois, M Massanella, N Brassard, N Alsahafi, GG Delgado, JP Routy, BD Walker, A Finzi, N Chomont, and DE Kaufmann. 2016. Single-Cell Characterization of Viral Translation-Competent Reservoirs in HIV-Infected Individuals. Cell Host Microbe 20: 368–380. https://doi.org/10.1016/j.chom.2016.07.015, PMID: 27545045

Bertagnolli LN, JA White, FR Simonetti, SA Beg, J Lai, C Tomescu, AJ Murray, AAR Antar, H Zhang, JB Margolick, R Hoh, SG Deeks, P Tebas, LJ Montaner, RF Siliciano, GM Laird, and JD Siliciano. 2018. The role of CD32 during HIV-1 infection. Nature 561: E17–e19. https://doi.org/10.1038/s41586-018-0494-3, PMID: 30232425

Boritz EA, S Darko, L Swaszek, G Wolf, D Wells, X Wu, AR Henry, F Laboune, J Hu, D Ambrozak, MS Hughes, R Hoh, JP Casazza, A Vostal, D Bunis, K Nganou-Makamdop, JS Lee, SA Migueles, RA Koup, M Connors, S Moir, T Schacker, F Maldarelli, SH Hughes, SG Deeks, and DC Douek. 2016. Multiple Origins of Virus Persistence during Natural Control of HIV Infection. Cell 166: 1004–1015. https://doi.org/10.1016/j.cell.2016.06.039, PMID: 27453467

Brenchley JM, BJ Hill, DR Ambrozak, DA Price, FJ Guenaga, JP Casazza, J Kuruppu, J Yazdani, SA Migueles, M Connors, M Roederer, DC Douek, and RA Koup. 2004. T-cell subsets that harbor human immunodeficiency virus (HIV) in vivo: implications for HIV pathogenesis. J Virol 78: 1160–8. https://doi.org/10.1128/jvi.78.3.1160-1168.2004, PMID: 14722271

Bruner KM, Z Wang, FR Simonetti, AM Bender, KJ Kwon, S Sengupta, EJ Fray, SA Beg, AAR Antar, KM Jenike, LN Bertagnolli, AA Capoferri, JT Kufera, A Timmons, C Nobles, J Gregg, N Wada, YC Ho, H Zhang, JB Margolick, JN Blankson, SG Deeks, FD Bushman, JD Siliciano, GM Laird, and RF Siliciano. 2019. A quantitative approach for measuring the reservoir of latent HIV-1 proviruses. Nature 566: 120–125. https://doi.org/10.1038/s41586-019-0898-8, PMID: 30700913

Cattin A, TR Wiche Salinas, A Gosselin, D Planas, B Shacklett, EA Cohen, MP Ghali, JP Routy, and P Ancuta. 2019. HIV-1 is rarely detected in blood and colon myeloid cells during viral-suppressive antiretroviral therapy. Aids 33: 1293–1306. https://doi.org/10.1097/qad.0000000000002195, PMID: 30870200

Chomont N, M El-Far, P Ancuta, L Trautmann, FA Procopio, B Yassine-Diab, G Boucher, MR Boulassel, G Ghattas, JM Brenchley, TW Schacker, BJ Hill, DC Douek, JP Routy, EK Haddad, and RP Sékaly. 2009. HIV reservoir size and persistence are driven by T cell survival and homeostatic proliferation. Nat Med 15: 893–900. https://doi.org/10.1038/nm.1972, PMID: 19543283

Cohn LB, IT da Silva, R Valieris, AS Huang, JCC Lorenzi, YZ Cohen, JA Pai, AL Butler, M Caskey, M Jankovic, and MC Nussenzweig. 2018. Clonal CD4(+) T cells in the HIV-1 latent reservoir display a distinct gene profile upon reactivation. Nat Med 24: 604–609. https://doi.org/10.1038/s41591-018-0017-7, PMID: 29686423

Darcis G, NA Kootstra, B Hooibrink, T van Montfort, I Maurer, K Groen, S Jurriaans, M Bakker, C van Lint, B Berkhout, and AO Pasternak. 2020. CD32(+)CD4(+) T Cells Are Highly Enriched for HIV DNA and Can Support Transcriptional Latency. Cell Rep 30: 2284–2296.e3. https://doi.org/10.1016/j.celrep.2020.01.071, PMID: 32075737

Deleage C, SW Wietgrefe, G Del Prete, DR Morcock, XP Hao, M Piatak, Jr., J Bess, JL Anderson, KE Perkey, C Reilly, JM McCune, AT Haase, JD Lifson, TW Schacker, and JD Estes. 2016. Defining HIV and SIV Reservoirs in Lymphoid Tissues. Pathog Immun 1: 68–106. https://doi.org/10.20411/pai.v1i1.100, PMID: 27430032

Deng W, BS Maust, DC Nickle, GH Learn, Y Liu, L Heath, SL Kosakovsky Pond, and JI Mullins. 2010. DIVEIN: a web server to analyze phylogenies, sequence divergence, diversity, and informative sites. Biotechniques 48: 405–8. https://doi.org/10.2144/000113370, PMID: 20569214

Descours B, G Petitjean, JL López-Zaragoza, T Bruel, R Raffel, C Psomas, J Reynes, C Lacabaratz, Y Levy, O Schwartz, JD Lelievre, and M Benkirane. 2017. CD32a is a marker of a CD4 T-cell HIV reservoir harbouring replication-competent proviruses. Nature 543: 564–567. https://doi.org/10.1038/nature21710, PMID: 28297712

Dudhane A, ZQ Wang, T Orlikowsky, A Gupta, GP Wormser, H Horowitz, P Kufer, and MK Hoffmann. 1996. AIDS patient monocytes target CD4 T cells for cellular conjugate formation and deletion through the membrane expression of HIV-1 envelope molecules. AIDS Res Hum Retroviruses 12: 893–9. https://doi.org/10.1089/aid.1996.12.893, PMID: 8798974

Durand CM, G Ghiaur, JD Siliciano, SA Rabi, EE Eisele, M Salgado, L Shan, JF Lai, H Zhang, J Margolick, RJ Jones, JE Gallant, RF Ambinder, and RF Siliciano. 2012. HIV-1 DNA is detected in bone marrow populations containing CD4+ T cells but is not found in purified CD34+ hematopoietic progenitor cells in most patients on antiretroviral therapy. J Infect Dis 205: 1014–8. https://doi.org/10.1093/infdis/jir884, PMID: 22275402

Fukazawa Y, R Lum, AA Okoye, H Park, K Matsuda, JY Bae, SI Hagen, R Shoemaker, C Deleage, C Lucero, D Morcock, T Swanson, AW Legasse, MK Axthelm, J Hesselgesser, R Geleziunas, VM Hirsch, PT Edlefsen, M Piatak, Jr., JD Estes, JD Lifson, and LJ Picker. 2015. B cell follicle sanctuary permits persistent productive simian immunodeficiency virus infection in elite controllers. Nat Med 21: 132–9. https://doi.org/10.1038/nm.3781, PMID: 25599132

Gosselin A, P Monteiro, N Chomont, F Diaz-Griffero, EA Said, S Fonseca, V Wacleche, M El-Far, MR Boulassel, JP Routy, RP Sekaly, and P Ancuta. 2010. Peripheral blood CCR4+CCR6+ and CXCR3+CCR6+CD4+ T cells are highly permissive to HIV-1 infection. J Immunol 184: 1604–16. https://doi.org/10.4049/jimmunol.0903058, PMID: 20042588

Grau-Expósito J, C Serra-Peinado, L Miguel, J Navarro, A Curran, J Burgos, I Ocaña, E Ribera, A Torrella, B Planas, R Badía, J Castellví, V Falcó, M Crespo, and MJ Buzon. 2017. A Novel Single-Cell FISH-Flow Assay Identifies Effector Memory CD4(+) T cells as a Major Niche for HIV-1 Transcription in HIV-Infected Patients. mBio 8. https://doi.org/10.1128/mBio.00876-17, PMID: 28698276

Green SA, M Smith, RB Hasley, D Stephany, A Harned, K Nagashima, S Abdullah, S Pittaluga, T Imamichi, J Qin, A Rupert, A Ober, HC Lane, and M Catalfamo. 2015. Activated platelet-T-cell conjugates in peripheral blood of patients with HIV infection: coupling coagulation/inflammation and T cells. Aids 29: 1297–308. https://doi.org/10.1097/qad.0000000000000701, PMID: 26002800

Günthard HF, SD Frost, AJ Leigh-Brown, CC Ignacio, K Kee, AS Perelson, CA Spina, DV Havlir, M Hezareh, DJ Looney, DD Richman, and JK Wong. 1999. Evolution of envelope sequences of human immunodeficiency virus type 1 in cellular reservoirs in the setting of potent antiviral therapy. J Virol 73: 9404–12. https://doi.org/10.1128/jvi.73.11.9404-9412.1999, PMID: 10516049

Ho YC, L Shan, NN Hosmane, J Wang, SB Laskey, DI Rosenbloom, J Lai, JN Blankson, JD Siliciano, and RF Siliciano. 2013. Replication-competent noninduced proviruses in the latent reservoir increase barrier to HIV-1 cure. Cell 155: 540–51. https://doi.org/10.1016/j.cell.2013.09.020, PMID: 24243014

Hogan LE, J Vasquez, KS Hobbs, E Hanhauser, B Aguilar-Rodriguez, R Hussien, C Thanh, EA Gibson, AB Carvidi, LCB Smith, S Khan, M Trapecar, S Sanjabi, M Somsouk, CA Stoddart, DR Kuritzkes, SG Deeks, and TJ Henrich. 2018. Increased HIV-1 transcriptional activity and infectious burden in peripheral blood and gut-associated CD4+ T cells expressing CD30. PLoS Pathog 14: e1006856. https://doi.org/10.1371/journal.ppat.1006856, PMID: 29470552

Iglesias-Ussel M, C Vandergeeten, L Marchionni, N Chomont, and F Romerio. 2013. High levels of CD2 expression identify HIV-1 latently infected resting memory CD4+ T cells in virally suppressed subjects. J Virol 87: 9148–58. https://doi.org/10.1128/jvi.01297-13, PMID: 23760244

Josefsson L, S Eriksson, E Sinclair, T Ho, M Killian, L Epling, W Shao, B Lewis, P Bacchetti, L Loeb, J Custer, L Poole, FM Hecht, and S Palmer. 2012. Hematopoietic precursor cells isolated from patients on long-term suppressive HIV therapy did not contain HIV-1 DNA. J Infect Dis 206: 28–34. https://doi.org/10.1093/infdis/jis301, PMID: 22536001

Josefsson L, S von Stockenstrom, NR Faria, E Sinclair, P Bacchetti, M Killian, L Epling, A Tan, T Ho, P Lemey, W Shao, PW Hunt, M Somsouk, W Wylie, DC Douek, L Loeb, J Custer, R Hoh, L Poole, SG Deeks, F Hecht, and S Palmer. 2013. The HIV-1 reservoir in eight patients on long-term suppressive antiretroviral therapy is stable with few genetic changes over time. Proc Natl Acad Sci U S A 110: E4987–96. https://doi.org/10.1073/pnas.1308313110, PMID: 24277811

Kearney MF, J Spindler, W Shao, S Yu, EM Anderson, A O’Shea, C Rehm, C Poethke, N Kovacs, JW Mellors, JM Coffin, and F Maldarelli. 2014. Lack of detectable HIV-1 molecular evolution during suppressive antiretroviral therapy. PLoS Pathog 10: e1004010. https://doi.org/10.1371/journal.ppat.1004010, PMID: 24651464

Kieffer TL, MM Finucane, RE Nettles, TC Quinn, KW Broman, SC Ray, D Persaud, and RF Siliciano. 2004. Genotypic analysis of HIV-1 drug resistance at the limit of detection: virus production without evolution in treated adults with undetectable HIV loads. J Infect Dis 189: 1452–65. https://doi.org/10.1086/382488, PMID: 15073683

Kuo HH, R Banga, GQ Lee, C Gao, M Cavassini, JM Corpataux, JE Blackmer, S Zur Wiesch, XG Yu, G Pantaleo, M Perreau, and M Lichterfeld. 2020. Blood and Lymph Node Dissemination of Clonal Genome-Intact Human Immunodeficiency Virus 1 DNA Sequences During Suppressive Antiretroviral Therapy. J Infect Dis 222: 655–660. https://doi.org/10.1093/infdis/jiaa137, PMID: 32236405

Lee GQ, N Orlova-Fink, K Einkauf, FZ Chowdhury, X Sun, S Harrington, HH Kuo, S Hua, HR Chen, Z Ouyang, K Reddy, K Dong, T Ndung’u, BD Walker, ES Rosenberg, XG Yu, and M Lichterfeld. 2017. Clonal expansion of genome-intact HIV-1 in functionally polarized Th1 CD4+ T cells. J Clin Invest 127: 2689–2696. https://doi.org/10.1172/jci93289, PMID: 28628034

Lorenzo-Redondo R, HR Fryer, T Bedford, EY Kim, J Archer, SLK Pond, YS Chung, S Penugonda, J Chipman, CV Fletcher, TW Schacker, MH Malim, A Rambaut, AT Haase, AR McLean, and SM Wolinsky. 2016. Persistent HIV-1 replication maintains the tissue reservoir during therapy. Nature 530: 51–56. https://doi.org/10.1038/nature16933, PMID: 26814962

Massanella M, W Bakeman, P Sithinamsuwan, JLK Fletcher, N Chomchey, S Tipsuk, T Chalermchai, JP Routy, J Ananworanich, VG Valcour, and N Chomont. 2019. Infrequent HIV Infection of Circulating Monocytes during Antiretroviral Therapy. J Virol 94. https://doi.org/10.1128/jvi.01174-19, PMID: 31597764

Mens H, AG Pedersen, LB Jørgensen, S Hue, Y Yang, J Gerstoft, and TL Katzenstein. 2007. Investigating signs of recent evolution in the pool of proviral HIV type 1 DNA during years of successful HAART. AIDS Res Hum Retroviruses 23: 107–15. https://doi.org/10.1089/aid.2006.0089, PMID: 17263640

Osuna CE, SY Lim, JL Kublin, R Apps, E Chen, TM Mota, SH Huang, Y Ren, ND Bachtel, AM Tsibris, ME Ackerman, RB Jones, DF Nixon, and JB Whitney. 2018. Evidence that CD32a does not mark the HIV-1 latent reservoir. Nature 561: E20–e28. https://doi.org/10.1038/s41586-018-0495-2, PMID: 30232424

Pardons M, AE Baxter, M Massanella, A Pagliuzza, R Fromentin, C Dufour, L Leyre, JP Routy, DE Kaufmann, and N Chomont. 2019. Single-cell characterization and quantification of translation-competent viral reservoirs in treated and untreated HIV infection. PLoS Pathog 15: e1007619. https://doi.org/10.1371/journal.ppat.1007619, PMID: 30811499

Pérez L, J Anderson, J Chipman, A Thorkelson, TW Chun, S Moir, AT Haase, DC Douek, TW Schacker, and EA Boritz. 2018. Conflicting evidence for HIV enrichment in CD32(+) CD4 T cells. Nature 561: E9–e16. https://doi.org/10.1038/s41586-018-0493-4, PMID: 30232423

Persaud D, GK Siberry, A Ahonkhai, J Kajdas, D Monie, N Hutton, DC Watson, TC Quinn, SC Ray, and RF Siliciano. 2004. Continued production of drug-sensitive human immunodeficiency virus type 1 in children on combination antiretroviral therapy who have undetectable viral loads. J Virol 78: 968–79. https://doi.org/10.1128/jvi.78.2.968-979.2004, PMID: 14694128

Raposo RAS, M de Mulder Rougvie, D Paquin-Proulx, PM Brailey, VD Cabido, PM Zdinak, AS Thomas, SH Huang, GA Beckerle, RB Jones, and DF Nixon. 2017. IFITM1 targets HIV-1 latently infected cells for antibody-dependent cytolysis. JCI Insight 2: e85811. https://doi.org/10.1172/jci.insight.85811, PMID: 28097226

Real F, C Capron, A Sennepin, R Arrigucci, A Zhu, G Sannier, J Zheng, L Xu, JM Massé, S Greffe, M Cazabat, M Donoso, P Delobel, J Izopet, E Eugenin, ML Gennaro, E Rouveix, E Cramer Bordé, and M Bomsel. 2020. Platelets from HIV-infected individuals on antiretroviral drug therapy with poor CD4(+) T cell recovery can harbor replication-competent HIV despite viral suppression. Sci Transl Med 12. https://doi.org/10.1126/scitranslmed.aat6263, PMID: 32188724

Ruff CT, SC Ray, P Kwon, R Zinn, A Pendleton, N Hutton, R Ashworth, S Gange, TC Quinn, RF Siliciano, and D Persaud. 2002. Persistence of wild-type virus and lack of temporal structure in the latent reservoir for human immunodeficiency virus type 1 in pediatric patients with extensive antiretroviral exposure. J Virol 76: 9481–92. https://doi.org/10.1128/jvi.76.18.9481-9492.2002, PMID: 12186930

Serra-Peinado C, J Grau-Expósito, L Luque-Ballesteros, A Astorga-Gamaza, J Navarro, J Gallego-Rodriguez, M Martin, A Curran, J Burgos, E Ribera, B Raventós, R Willekens, A Torrella, B Planas, R Badía, F Garcia, J Castellví, M Genescà, V Falcó, and MJ Buzon. 2019. Expression of CD20 after viral reactivation renders HIV-reservoir cells susceptible to Rituximab. Nat Commun 10: 3705. https://doi.org/10.1038/s41467-019-11556-4, PMID: 31420544

Simpson SR, MV Singh, S Dewhurst, G Schifitto, and SB Maggirwar. 2020. Platelets function as an acute viral reservoir during HIV-1 infection by harboring virus and T-cell complex formation. Blood Adv 4: 4512–4521. https://doi.org/10.1182/bloodadvances.2020002420, PMID: 32946568

Telwatte S, S Lee, M Somsouk, H Hatano, C Baker, P Kaiser, P Kim, TH Chen, J Milush, PW Hunt, SG Deeks, JK Wong, and SA Yukl. 2018. Gut and blood differ in constitutive blocks to HIV transcription, suggesting tissue-specific differences in the mechanisms that govern HIV latency. PLoS Pathog 14: e1007357. https://doi.org/10.1371/journal.ppat.1007357, PMID: 30440043

Thornhill JP, M Pace, GE Martin, J Hoare, S Peake, C Herrera, C Phetsouphanh, J Meyerowitz, E Hopkins, H Brown, P Dunn, N Olejniczak, C Willberg, P Klenerman, R Goldin, J Fox, S Fidler, and J Frater. 2019. CD32 expressing doublets in HIV-infected gut-associated lymphoid tissue are associated with a T follicular helper cell phenotype. Mucosal Immunol 12: 1212–1219. https://doi.org/10.1038/s41385-019-0180-2, PMID: 31239514

Vásquez JJ, BL Aguilar-Rodriguez, L Rodriguez, LE Hogan, M Somsouk, JM McCune, SG Deeks, ZG Laszik, PW Hunt, and TJ Henrich. 2019. CD32-RNA Co-localizes with HIV-RNA in CD3+ Cells Found within Gut Tissues from Viremic and ART-Suppressed Individuals. Pathog Immun 4: 147–160. https://doi.org/10.20411/pai.v4i1.271, PMID: 31139759

Yukl SA, S Gianella, E Sinclair, L Epling, Q Li, L Duan, AL Choi, V Girling, T Ho, P Li, K Fujimoto, H Lampiris, CB Hare, M Pandori, AT Haase, HF Günthard, M Fischer, AK Shergill, K McQuaid, DV Havlir, and JK Wong. 2010. Differences in HIV burden and immune activation within the gut of HIV-positive patients receiving suppressive antiretroviral therapy. J Infect Dis 202: 1553–61. https://doi.org/10.1086/656722, PMID: 20939732

